# EXPRESSION OF A SECRETABLE, CELL-PENETRATING CDKL5 PROTEIN ENHANCES THE EFFICACY OF AAV VECTOR-MEDIATED GENE THERAPY FOR CDKL5 DEFICIENCY DISORDER

**DOI:** 10.1101/2021.07.26.453746

**Authors:** Giorgio Medici, Marianna Tassinari, Giuseppe Galvani, Stefano Bastianini, Laura Gennaccaro, Manuela Loi, Nicola Mottolese, Sara Alvente, Chiara Berteotti, Giulia Sagona, Leonardo Lupori, Helen Rappe Baggett, Giovanna Zoccoli, Maurizio Giustetto, Alysson Muotri, Tommaso Pizzorusso, Hiroyuki Nakai, Stefania Trazzi, Elisabetta Ciani

**Affiliations:** Department of Biomedical and Neuromotor Science, University of Bologna, Bologna, Italy; Department of Developmental Neuroscience, IRCCS Stella Maris Foundation, 56128 Pisa, Italy. Department of Neuroscience, Psychology, Drug Research and Child Health (NEUROFARBA), University of Florence, Florence, Italy; Department of Developmental Neuroscience, IRCCS Stella Maris Foundation, Pisa, Italy. Scuola Normale Superiore, Pisa, Italy; Departments of Molecular and Medical Genetics and Molecular Immunology and Microbiology, Oregon Health & Science University, Portland, OR 97239, USA; Department of Neuroscience “Rita Levi Montalcini”, University of Turin, Turin, Italy; University of California San Diego, School of Medicine, Department of Pediatrics/Rady Children’s Hospital-San Diego, Department of Cellular & Molecular Medicine, Kavli Institute for Brain and Mind, Archealization Center (ArchC), Center for Academic Research and Training in Anthropogeny (CARTA), La Jolla, CA 92037, USA; Institute of Neuroscience, National Research Council, 56124 Pisa, Italy; Scuola Normale Superiore, Pisa, Italy; Division of Neuroscience, Oregon National Primate Research Center, Beaverton, OR 97006, USA

## Abstract

No therapy is currently available for CDKL5 (cyclin-dependent kinase-like 5) deficiency disorder (CDD), a severe neurodevelopmental disorder caused by mutations in the *CDKL5* gene. Although delivery of a wild-type copy of the mutated gene to cells represents the most curative approach for a monogenic disease, proof-of-concept studies highlight significant efficacy caveats for brain gene therapy. Herein, we used a secretable TATk-CDKL5 protein to enhance the efficiency of a gene therapy for CDD. We found that, although AAVPHP.B_Igk-TATk-CDKL5 and AAVPHP.B_CDKL5 vectors had similar brain infection efficiency, the AAVPHP.B_Igk-TATk-CDKL5 vector led to a higher CDKL5 protein replacement due to secretion and transduction of the TATk-CDKL5 protein into the neighboring cells. Importantly, *Cdkl5* KO mice treated with the AAVPHP.B_Igk-TATk-CDKL5 vector showed a behavioral and neuroanatomical improvement in comparison with vehicle-treated *Cdkl5* KO mice or *Cdkl5* KO mice treated with the AAVPHP.B_CDKL5 vector, indicating that a gene therapy based on a secretable recombinant TATk-CDKL5 protein is more effective at compensating *Cdkl5*-null brain defects than gene therapy based on the expression of the native CDKL5.

## INTRODUCTION

CDKL5 (cyclin-dependent kinase-like 5) deficiency disorder (CDD) is a severe X-linked neurodevelopmental disease caused by mutations in the *CDKL5* gene, which lead to a lack of CDKL5 protein expression or function. CDD mainly affects girls and is characterized by early-onset epileptic seizures, hypotonia, intellectual disability, motor and visual impairment, and, in some cases, respiratory dysregulation ^1–6^. Although pharmacological treatments are used to control seizures, there is currently no cure or effective treatment to ameliorate cognitive and behavioral symptoms for CDD. Animal models of CDKL5 disorder, *Cdkl5* knockout (KO) mice ^7–9^, recapitulate different features of CDD, exhibiting severe impairment in learning and memory, visual and respiratory deficits, and motor stereotypies ^7,8,10–12^ and, therefore, they are a good model with which to study the positive effects of therapeutic strategies.

In theory, for a monogenic disease such as CDD, the delivery of a wild-type copy of the mutated gene to cells which lack functional protein represents the most curative approach. Using AAV, advancements in global central nervous system (CNS) gene delivery have accelerated to the point that treatments for neurodevelopmental disorders, such as lysosomal storage disease, Rett Syndrome, CDD, Fragile X, and autism seem within reach ^13^. However, gene therapy is not without risks for humans. The major caveat regards the low efficiency of gene delivery to the CNS by viral vectors that requires large vector doses, and consequently, brings the risk of immune reaction, as was the case in human clinical trials for hemophilia B ^14,15^. Moreover, the new gene might be inserted into the DNA in the wrong location, possibly causing harmful mutations to the DNA or even cancer, as shown in rodent studies ^16^. If the protein produced by the viral vector-infected cells can be secreted and enter into neighboring cells, this will amplify the effect of the gene therapy because, even if the transduced cells are low in number, they will become a “factory” for the production of the therapeutic protein, supplying therapeutic molecules to neighboring cells. In this scenario, the efficiency of gene delivery does not necessarily need to be high. This decreases the risk of insertional mutagenesis and toxic side effects connected with large vector doses.

We recently created an Igk-TATk-CDKL5 fusion construct and found that, in view of the properties of the Igk-chain leader sequence, the TATk-CDKL5 protein produced by infected cells is secreted via constitutive secretory pathways ^17^. Importantly, due to the transduction property of the TATk peptide, the secreted CDKL5 protein is internalized by cells, maintaining its native biological activity^17^.

Here we compared the effects of a CDKL5 gene therapy with an IgK-TATk-CDKL5 gene therapy in a *Cdkl5* KO mouse model to validate whether the Igk-TATk-CDKL5 approach, that enhances the biodistribution of the therapeutic CDKL5 kinase from genetically-corrected cells to non-corrected cells via a cross-correction mechanism, significantly enhances the therapeutic efficacy.

## RESULTS

### Production of an AAVPHP.B_Igk-TATk-CDKL5 vector for the expression of a secretable TATk-CDKL5 protein

To express the Igk-TATk-CDKL5 and CDKL5 proteins by means of AAV vector-mediated gene delivery, an Igk-TATk-CDKL5 or a CDKL5 gene expression cassette ^17^ was inserted into the AAV vector plasmid under the control of a strong, long-term, and ubiquitous expression promoter ^18^. The efficiency of the newly-generated AAV vectors to express CDKL5 and Igk-TATk-CDKL5 proteins in neurons was confirmed after primary hippocampal neuron infection (Fig. 1A,B). Igk-TATk-CDKL5 showed a different expression pattern compared to CDKL5, with a more cytoplasmic distribution (Fig. 1A,B) that suggests the presence of protein secretion via the constitutive secretory pathways. We confirmed efficient TATk-CDKL5 protein secretion using western blot analysis (Fig. 1C). TATk-CDKL5, but not CDKL5 protein, was detected in the culture medium of HEK293T cells transfected with the AAVPHP.B_Igk-TATk-CDKL5 or the AAVPHP.B_CDKL5 vector plasmid DNA (Fig. 1C). To investigate whether neurons are transduced by secreted TATk-CDKL5 protein, hippocampal neurons were co-cultured, using transwells (Fig. 1D), with HEK293T cells transfected with the AAV-Igk-TATk-CDKL5 or the AAV-CDKL5 plasmid. After 48 h of co-culture, TATk-CDKL5 protein was efficiently internalized by hippocampal neurons (Fig. 1D), while, as expected, no CDKL5-positive neurons were present (data not shown).

**Figure 1.**
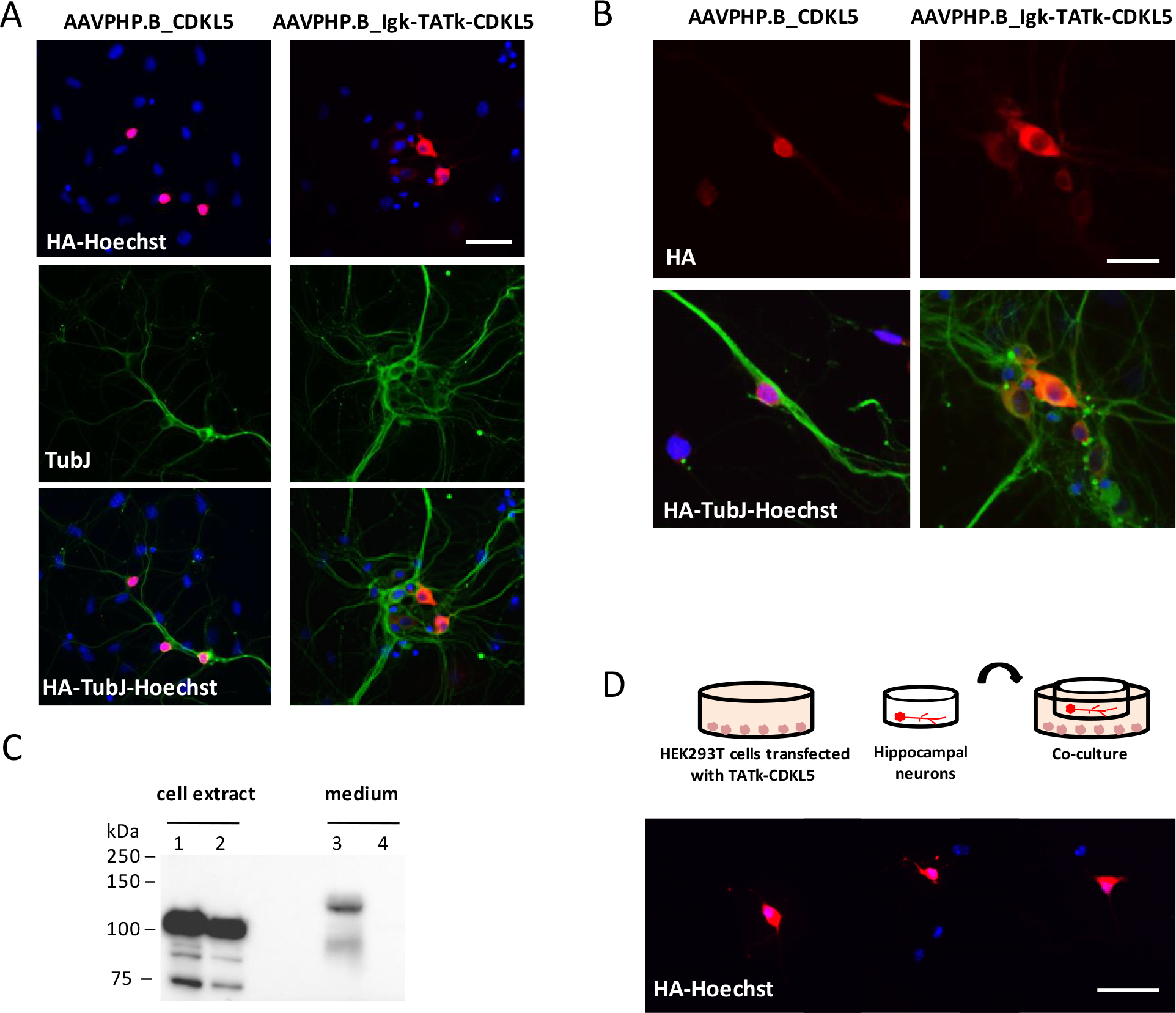
Secretion and transduction efficiency of the TATk-CDKL5 protein. (A, B) Two-day (DIV2) primary hippocampal neuronal cultures from *Cdkl5* −/Y mice were infected with the AAVPHP.B_Igk-TATk-CDKL5 or the AAVPHP.B_CDKL5 vector (MOI 10^6^) and fixed at DIV 7. CDKL5 and TATk-CDKL5 protein localization was assessed by immunostaining with an anti-HA antibody (red) and an anti-β III tubulin antibody (TubJ, green). Nuclei were counterstained with Hoechst. Scale bars = 50 µ m (A), 25 μm (B). (C) Western blot analysis using an anti-HA antibody confirmed TATk-CDKL5 and CDKL5 protein expression in AAV vector plasmid DNA-transfected HEK293T cells (cell extract, lanes 1 and 2), and TATk-CDKL5 protein accumulation in the concentrated culture medium (lane 3), indicating that the TATk-CDKL5 protein was secreted from cells. No CDKL5 expression was detected in the medium of HEK293T cells transfected with the AAVPHP.B_CDKL5 vector (lane 4). (D) Co-culture experimental design: HEK293T cells were transfected with the AAV vector plasmid containing the Igk-TATk-CDKL5 cassette. Twenty-four hours after transfection, cover glasses with 5-day (DIV5) differentiated primary hippocampal neurons were transferred to the HEK293T 6-well plate in an elevated position. Fluorescence microscopy images showing the presence of TATk-CDKL5 protein in differentiated primary hippocampal neurons from *Cdkl5* −/Y mice co-cultured for 48 h (DIV5-DIV7) with HEK293T cells transfected with AAVPHP.B_Igk-TATk-CDKL5 plasmid. Neurons were immunostained with an anti-HA antibody (red) and nuclei were counterstained with Hoechst. Scale bar = 50 µ m.

The efficiency of TATk-CDKL5 protein transduction in vivo was analysed in *Cdkl5* KO (−/Y) mice that had undergone intraventricular injection with AAVPHP.B_Igk-TATk-CDKL5 vector at the neonatal stage and that were sacrificed 2 months after the injection. By combining fluorescent in situ hybridization (ISH) and immunohistochemical staining to simultaneously visualize CDKL5 mRNA and protein, respectively, we found CDKL5 protein replacement in brain cells that did not show ISH staining (Fig. 2), indicating a cross-correction mechanism mediated by the secretable, cell-penetrating TATk-CDKL5 protein.

**Figure 2.**
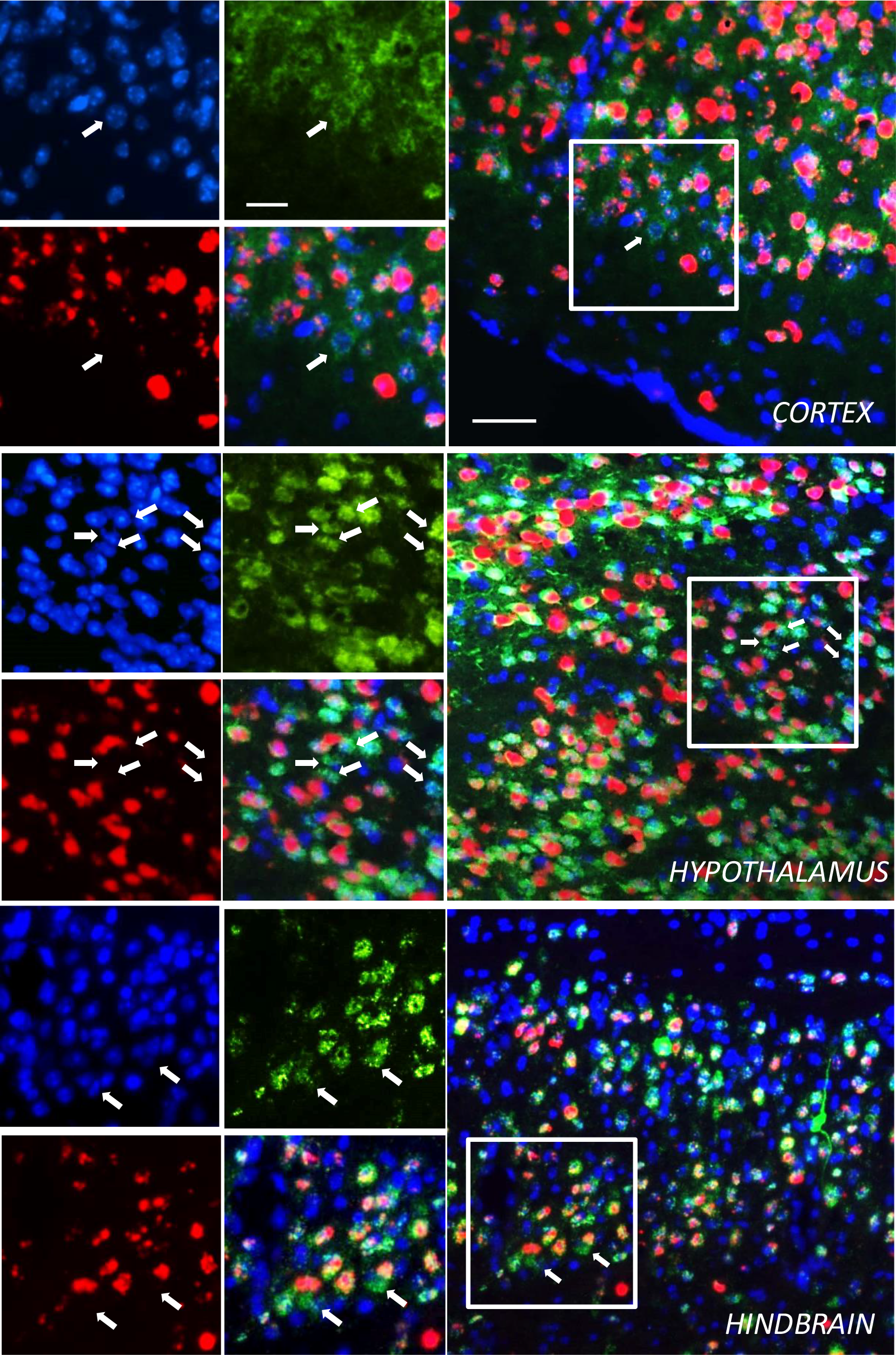
*TATk-CDKL5* mRNA and protein distribution in mouse brain sections. Fluorescence *In Situ* Hybridization (ISH) for *CDKL5* mRNA combined with fluorescence immunolabeling for TATk-CDKL5 protein in mouse brain sections of 2-month-old *Cdkl5* −/Y mice intraventricularly injected at the neonatal stage with AAVPHP.B_Igk-TATk-CDKL5 vector. Images show *TATk-CDKL5* mRNA (red) and protein (green) localization in the cortex, hypothalamus, and hindbrain of a treated mouse 90 days post-injection. Localization of *TATk-CDKL5* mRNA was evaluated through ISH with a CDKL5 probe, while TATk-CDKL5 protein was evaluated through immunohistochemistry using an anti-HA antibody; nuclei were counterstained with DAPI. The white boxes indicate the regions shown in the high magnification panels. The white arrows indicate cross-corrected cells (HA-immunopositive cells with no ISH staining). Scale bar = 50 µ m (low magnification); 25 µ m (high magnification).

### Effect of gene therapy with AAVPHP.B_Igk-TATk-CDKL5 or AAVPHP.B_CDKL5 vector on behavior in C*dkl5* −/Y mice

To evaluate the effectiveness of a cross-correction mechanism compared to a classic gene therapy approach, adult (3-4 months old) *Cdkl5* −/Y mice were administered with AAVPHP.B_CDKL5 or AAVPHP.B_Igk-TATk-CDKL5 vector at a dose of 10^12^ vg/mouse via intracarotid injection, and the effects of treatment were evaluated 60 days post-injection (Supplementary Fig. 1A). A group of vehicle-treated *Cdkl5* −/Y and wild-type (+/Y) mice were used as controls for behavioral tests. Importantly, no changes in terms of body weight, sleep pattern, or microglial cell number were observed in vector-treated *Cdkl5* −/Y mice compared to age-matched vehicle-treated mice (Supplementary Fig. 1B-D), indicating that viral infection and secreted CDKL5 protein did not affect animal well-being and/or cause an inflammatory response.

Loss of Cdkl5 function in *Cdkl5* −/Y mice is associated with autistic-like (ASD-like) phenotypes, analysed through home-cage social behaviors (marble burying and nest building ability) ^19^. *Cdkl5* −/Y mice buried a significantly lower number of marbles and showed a reduced nest building ability compared to wild-type (+/Y) mice (Fig. 3A,B). Sixty days after treatment with AAVPHP.B_Igk-TATk-CDKL5 vector *Cdkl5* −/Y mice buried a higher number of marbles compared to vehicle-treated and AAVPHP.B_CDKL5-treated *Cdkl5* −/Y mice (Fig. 3A). Similarly, nest building ability was improved only in *Cdkl5* −/Y mice treated with AAVPHP.B_Igk-TATk-CDKL5 vector (Fig. 3B).

**Figure 3.**
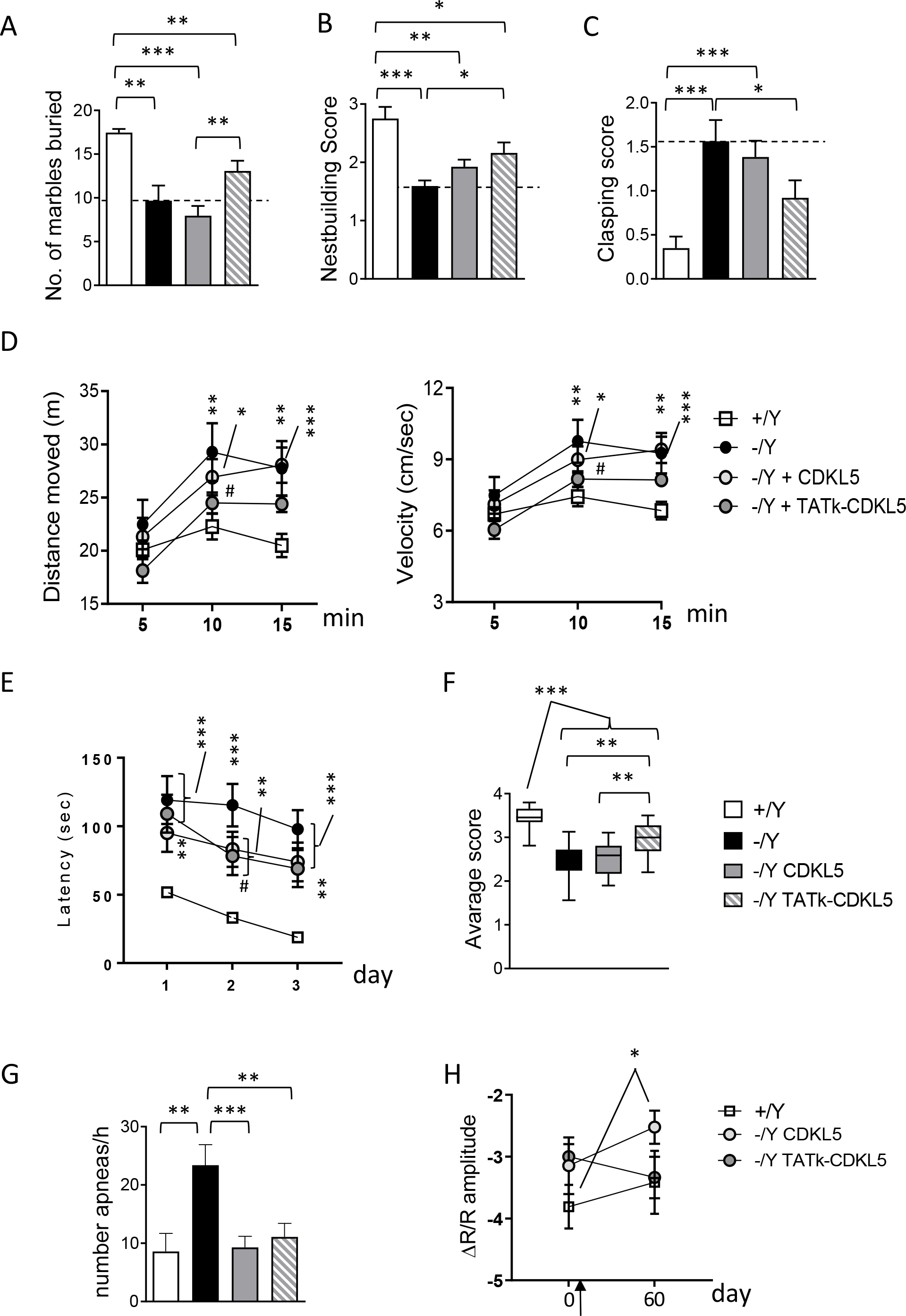
Effect of CDKL5 and TATk-CDKL5 proteins on behavior in *Cdkl5* −/Y mice. (A, B) Autistic-like features in treated *Cdkl5* −/Y mice. Number of marbles buried (A) and nest quality (B) of wild-type mice (+/Y, n = 20) and *Cdkl5* −/Y mice (n = 6 (A); n = 16 (B)) and of *Cdkl5* −/Y mice 60 days from treatment with AAVPHP.B_CDKL5 (n = 26) or AAVPHP.B_Igk-TATk-CDKL5 (n = 25). (C) Hind-limb clasping score during a 10 sec interval in wild-type mice (+/Y, n = 20) and *Cdkl5* −/Y mice (n = 16), and in *Cdkl5* −/Y mice 60 days after treatment with AAVPHP.B_CDKL5 (n = 26) or AAVPHP.B_Igk-TATk-CDKL5 (n = 25). (D) Total distance traveled (left graph) and average locomotion velocity (right graph) of wild-type mice (+/Y, n = 20) and *Cdkl5* −/Y mice (n = 16), and of *Cdkl5* −/Y mice treated with AAVPHP.B_CDKL5 (n = 26) or AAVPHP.B_Igk-TATk-CDKL5 (n = 25), during a 15 min open field test. (E) Spatial learning was assessed using the Barnes Maze in wild-type mice (+/Y, n = 16) and *Cdkl5* −/Y mice (n = 10), and in *Cdkl5* −/Y mice treated with AAVPHP.B_CDKL5 (n = 15) or AAVPHP.B_Igk-TATk-CDKL5 (n = 13). Graphs show the mean latency to find the target hole during the 3-day learning period. (F) Behavioral score was assessed on a 1-4 scale for marble burying, nesting test, hindlimb clasping, open field (velocity and distance moved), and Barnes maze tests. The average score for each genotype and treatment was calculated. Whiskers show minimum and maximum score values among the same group of animals. **p < 0.01; ***p < 0.001 (Dunn’s test after Kruskall-Wallis). (G) Sleep apnea occurrence rate in treated *Cdkl5* −/Y mice was assessed using whole-body plethysmography. Sleep apnea occurrence in vehicle-treated wild-type (+/Y, n = 9) and *Cdkl5* −/Y (n = 7) mice, and in *Cdkl5* −/Y mice treated with AAVPHP.B_CDKL5 (n = 16) or AAVPHP.B_Igk-TATk-CDKL5 (n = 15), during rapid eye movement sleep (REMS). (H) Mean amplitude of visually evoked IOS responses measured before and after 60 days in vehicle-treated (n = 7) *Cdkl5* +/Y mice and AAVPHP.B_CDKL5 (n = 8) or AAVPHP.B_Igk-TATk-CDKL5 (n = 6) treated *Cdkl5* −/Y. The black arrow indicates the treatment time, one week after the first visually evoked IOS response measurement. Values (A, B, C, G) are presented as means ± SE. *p < 0.05; **p < 0.01; ***p < 0.001 (datasets in A, B, and C, Dunn’s test after Kruskall-Wallis; datasets in G, Fisher’s LSD test after one-way ANOVA). Values in (D, E, H) are presented as means ± SE. **p < 0.01; ***p < 0.001 compared to the vehicle-treated wild-type condition; #p < 0.05 as compared to the vehicle-treated *Cdkl5* −/Y samples (Fisher’s LSD test after repeated two-way ANOVA).

Stereotypic movements characterize *Cdkl5* −/Y mice ^8,19^ and CDD patients ^20^. In order to examine the effect of gene therapy on motor stereotypies, mice were tested for hindlimb clasping (Fig. 3C). Unlike wild-type (+/Y) mice, vehicle-treated and AAVPHP.B_CDKL5-treated *Cdkl5* −/Y mice showed a higher clasping score (Fig. 3C). On the contrary, *Cdkl5* −/Y mice treated with AAVPHP.B_Igk-TATk-CDKL5 vector showed a decrease in clasping (Fig. 3C), indicating that gene therapy with Igk-TATk-CDKL5 has a greater positive impact on the stereotypic behavior that is due to loss of Cdkl5 expression.

We assessed motor function of treated *Cdkl5* −/Y mice in the open-field test. The elevated locomotor activity (longer distance traveled with a higher average speed; Fig. 3D) that characterizes *Cdkl5* −/Y mice was improved by treatment with AAVPHP.B_Igk-TATk-CDKL5 vector (Fig. 3D), while no improvement was observed in mice treated with AAVPHP.B_CDKL5 vector.

Learning and memory were evaluated using the Barnes maze test, a cognitive paradigm in which *Cdkl5* −/Y mice are documented to be impaired ^21^. A significative improvement in learning was observed only in *Cdkl5* −/Y mice treated with AAVPHP.B_Igk-TATk-CDKL5 vector (Fig. 3E). Differently, no memory improvement was observed in *Cdkl5* −/Y mice treated with AAVPHP.B_Igk-TATk-CDKL5 or AAVPHP.B_CDKL5 vector during the probe trial on the 4th day (Supplementary Fig. 2A).

An overall mean phenotype score analysis including ASD-like phenotypes (Fig. 3A,B), stereotypic movements (Fig. 3C), motor function (Fig. 3D), and learning (Fig. 3E) showed a significative improvement only in *Cdkl5* −/Y mice treated with AAVPHP.B_Igk-TATk-CDKL5 vector (Fig. 3F), indicating that gene therapy with Igk-TATk-CDKL5 has a greater positive impact on behavior.

### Effect of gene therapy with AAVPHP.B_Igk-TATk-CDKL5 or AAVPHP.B_CDKL5 vector on breathing pattern and cortical visual responses in *Cdkl5* −/Y mice

Since impaired breathing pattern, particularly during sleep, and visual responses represent promising biomarkers for preclinical and clinical studies on CDD ^17,19,22,23^, we evaluated the effect of the treatments with AAVPHP.B_Igk-TATk-CDKL5 and AAVPHP.B_CDKL5 vector on these two patterns in *Cdkl5* −/Y mice. Using whole-body plethysmography, we found that treatment with both AAVPHP.B_CDKL5 and AAVPHP.B_Igk-TATk-CDKL5 vectors led to a drastic reduction in the number of apneas during REM sleep that became similar to that of wild-type mice (Fig. 3G). In contrast, an improvement was not achieved during NREM sleep (Supplementary Fig. 2B).

Cortical visual responses were assessed using non-invasive transcranial intrinsic optical signal (IOS) imaging, a method that allows us to monitor the visually evoked responses in the same animal at different time points ^23^. We found that, while AAVPHP.B_CDKL5-treated *Cdkl5* ™/Y mice showed a significantly reduced response with respect to the wild-type baseline condition (Fig. 3H), the visual responses of *Cdkl5* ™/Y mice treated with the AAVPHP.B_Igk-TATk-CDKL5 vector underwent an improvement, becoming similar to those found in wild-type mice (Fig. 3H).

### Effect of gene therapy with AAVPHP.B_Igk-TATk-CDKL5 or AAVPHP.B_CDKL5 vector on dendritic hypotrophy and connectivity in the hippocampus of *Cdkl5* −/Y mice

Dendritic arborization was found to be reduced in cortical and hippocampal pyramidal neurons of *Cdkl5* −/Y mice ^8,11,24,25^. In addition, *Cdkl5* −/Y mice exhibit a deficit in dendritic spine structure and stabilization ^11,24–27^ and a reduction in the number of PSD-95-positive puncta ^17,25^, which indicates loss of excitatory synaptic contacts. In order to establish the effect of gene therapy on dendritic pattern, we evaluated the dendritic length and spine density of CA1 pyramidal neurons (Fig. 4). In *Cdkl5* −/Y mice treated with AAVPHP.B_Igk-TATk-CDKL5 vector the length of both apical and basal dendrites was recovered compared to vehicle-treated and AAVPHP.B_CDKL5-treated *Cdkl5* −/Y mice (Fig. 4A,C), an improvement which may be mainly attributable to a recovery in the number of branches (Fig. 4B,C). Unlike in AAVPHP.B_CDKL5-treated *Cdkl5* −/Y mice, we found a recovery of spine density in *Cdkl5* −/Y mice treated with AAVPHP.B_Igk-TATk-CDKL5 vector in comparison with vehicle-treated *Cdkl5* −/Y mice (Fig. 4D,F). The percentage of mature spines was restored in *Cdkl5* −/Y mice treated with AAVPHP.B_Igk-TATk-CDKL5 vector (Fig. 4E), and it was only partially recovered in *Cdkl5* −/Y mice treated with AAVPHP.B_CDKL5 (Fig. 4E). Similarly, the number of PSD-95 puncta was restored in AAVPHP.B_Igk-TATk-CDKL5 treated *Cdkl5* −/Y mice (Fig. 4G,H), while only a partial improvement was present in *Cdkl5* −/Y mice treated with AAVPHP.B_CDKL5 (Fig. 4G,H).

**Figure 4.**
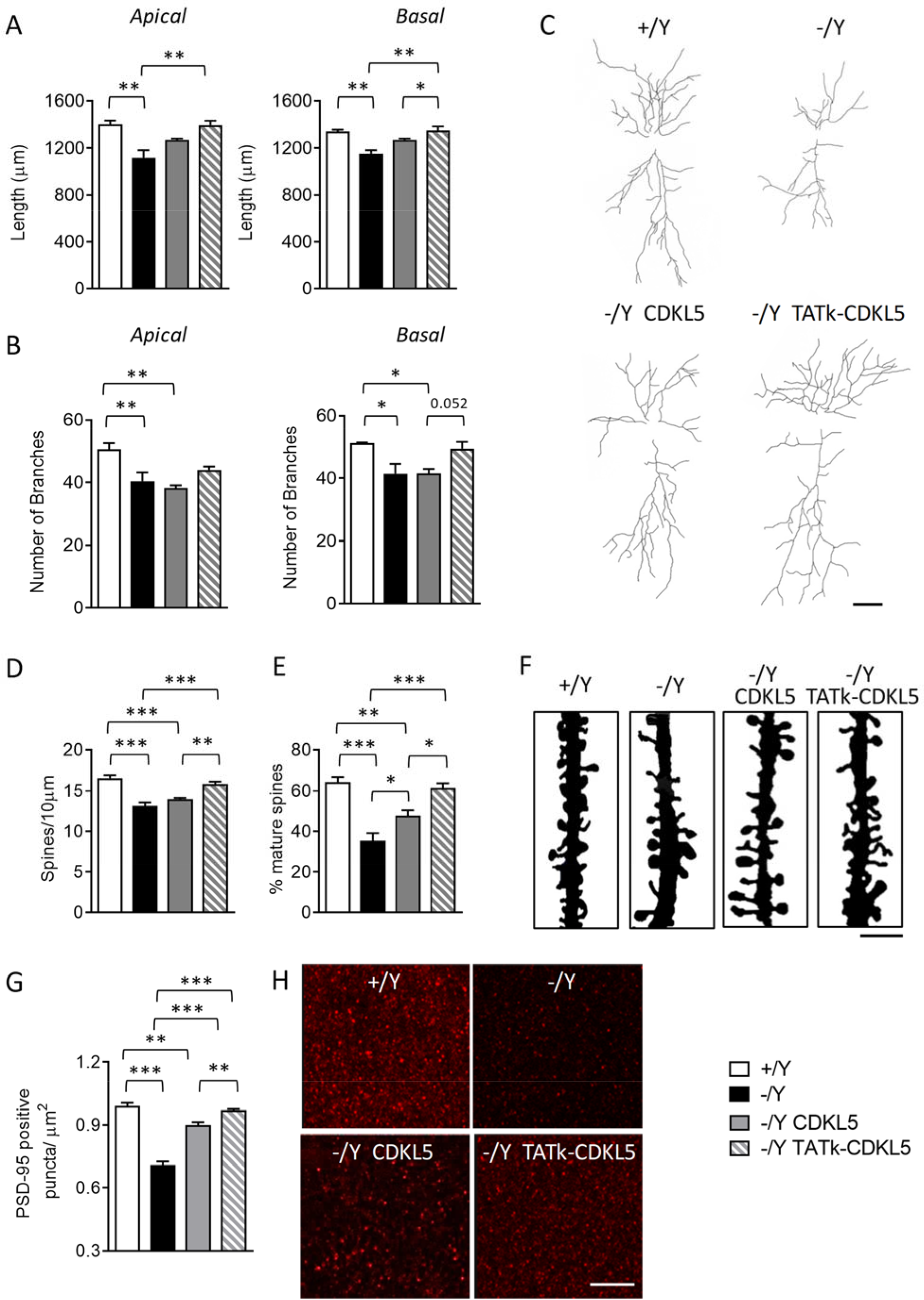
Effect of CDKL5 and TATk-CDKL5 proteins on dendritic morphology and connectivity. (A,B) Apical and basal mean total dendritic length (A) and mean number of dendritic segments (B) of Golgi-stained CA1 pyramidal neurons of wild-type (+/Y, n = 3) and *Cdkl5* −/Y (n = 3) mice and of *Cdkl5* −/Y mice 90 days from treatment with AAVPHP.B_CDKL5 (n = 4) or AAVPHP.B_Igk-TATk-CDKL5 (n = 4). (C) Example of the reconstructed apical and basal dendritic tree of Golgi-stained CA1 pyramidal neurons of 1 animal from each experimental group. Scale bar = 50 µ m (D, E) Dendritic spine density (D) and percentage of mature spines in relation to the total number of protrusions (E) of CA1 pyramidal neurons from wild-type (+/Y, n = 3) and *Cdkl5* −/Y (n = 3) mice and of *Cdkl5* −/Y mice treated with AAVPHP.B_CDKL5 (n = 4) or AAVPHP.B_Igk-TATk-CDKL5 (n = 4). (F) Images of Golgi-stained dendritic branches of CA1 pyramidal neurons of 1 animal from each experimental group. Scale bar = 2 µ m. (G) Number of fluorescent puncta per µ m^2^ exhibiting PSD-95 immunoreactivity in the CA1 layer of the hippocampus of wild-type (+/Y, n = 3) and *Cdkl5* −/Y (n = 3) mice and of *Cdkl5* −/Y mice treated with AAVPHP.B_CDKL5 (n = 4) or AAVPHP.B_Igk-TATk-CDKL5 (n = 4). (H) Representative fluorescence image of PSD-95 immunoreactive puncta in the hippocampus of 1 animal from each experimental group. Scale bar = 6 μm. Values are represented as means ± SE. *P< 0.05; **P < 0.01; ***P < 0.001 (datasets in A, B, D and G, Turkey’s test after one-way ANOVA; dataset in E, Dunn’s test after a Kruskal-Wallis test).

### Effect of gene therapy with AAVPHP.B_Igk-TATk-CDKL5 or AAVPHP.B_CDKL5 vector on neuronal survival and microglia activation in *Cdkl5* −/Y mice

*Cdkl5* −/Y mice are characterized by decreased survival of hippocampal neurons ^28,29^, and by an increased microglial activation ^30^. *Cdkl5* −/Y mice treated with AAVPHP.B_Igk-TATk-CDKL5 vector showed a higher number of Hoechst-positive nuclei and NeuN-positive pyramidal neurons in the CA1 layer (Fig. 5A-C) in comparison with vehicle-or AAVPHP.B_CDKL5-treated *Cdkl5* −/Y mice, indicating that a gene therapy with Igk-TATk-CDKL5 has a greater positive impact on the impaired neuronal survival that is due to loss of Cdkl5 expression. Similarly, a reversal of the inflammatory status, with a reduction in microglial soma size compared to the control levels, was present only in *Cdkl5* −/Y mice treated with AAVPHP.B_Igk-TATk-CDKL5 vector (Fig. 5D,E).

**Figure 5.**
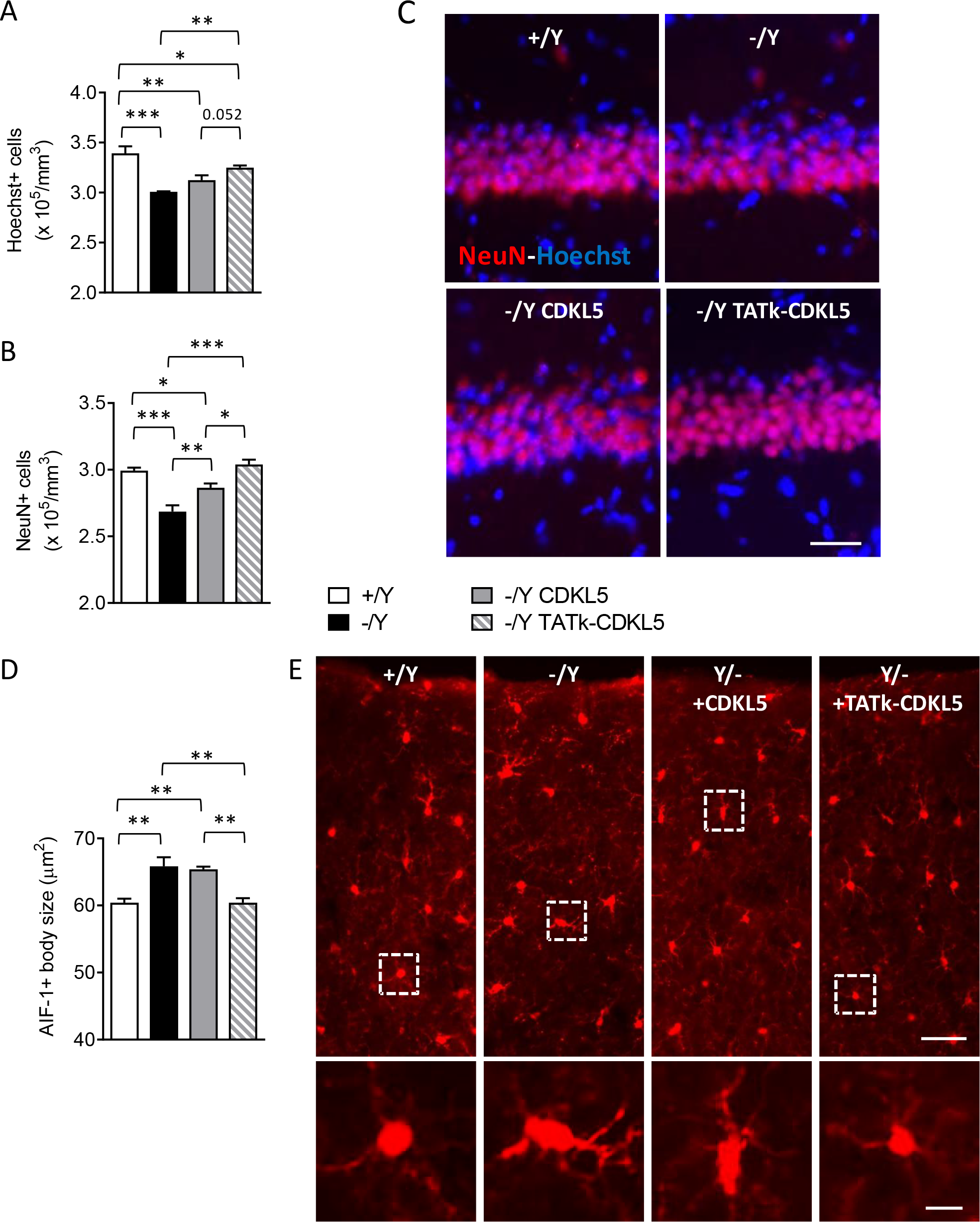
Effect of CDKL5 and TATk-CDKL5 proteins on neuronal survival and microglia activation in the brain of *Cdkl5* −/Y mice. (A, B) Quantification of Hoechst-positive cells (A) and NeuN-positive cells (B) in the CA1 layer of hippocampal sections from wild-type (+/Y, n = 3) and *Cdkl5* −/Y (n = 3) mice and from *Cdkl5* −/Y mice treated with AAVPHP.B_CDKL5 (n = 4) or AAVPHP.B_Igk-TATk-CDKL5 (n = 4). (C) Representative fluorescence images of sections that were immunopositive for NeuN (red) and counterstained with Hoechst (blue) in the hippocampal CA1 region of 1 animal from each group. Scale bar = 50 μm. (D) Mean microglia cell body size in cortical sections from wild-type (+/Y, n = 3) and *Cdkl5* −/Y (n = 3) mice, and from *Cdkl5* −/Y mice treated with AAVPHP.B_CDKL5 (n = 4) or AAVPHP.B_Igk-TATk-CDKL5 (n = 4). (E) Representative fluorescence images of cortical sections processed for AIF-1 immunohistochemistry of one animal from each group. The dotted boxes in the upper panels indicate microglial cells shown in magnification in the lower panels. Scale bar = 50 μm (low magnification), 10 μm (high magnification). Values are presented as means ± SE. *p < 0.05; **p < 0.01; ***p < 0.001 (Fisher’s LSD test after one-way ANOVA).

### Evaluation of the efficiency of AAV vector transduction and CDKL5 protein biodistribution

To quantify and compare the efficiency of gene transfer between AAVPHP.B_Igk-TATk-CDKL5 and AAVPHP.B_CDKL5 vectors, we assessed vector genome copy numbers per cell in several brain regions via qPCR. We found that AAVPHP.B_Igk-TATk-CDKL5 and AAVPHP.B_CDKL5 vectors had the same brain transduction efficiency, regardless of the tropism of various brain regions (Fig. 6A) and induced similar *CDKL5* mRNA levels in the brains of treated mice (Fig. 6B, Supplementary Fig. 3A). By comparing the *CDKL5* mRNA levels in treated *Cdkl5* −/Y mice to those of wild-type mice we found that the levels in treated mice were lower than those of wild-type mice in the cortex and hippocampus (Fig. 6B, Supplementary Fig. 3A), indicating only a partial recovery of CDKL5 expression in these brain regions in *Cdkl5* −/Y mice. Only in the hindbrain did the *CDKL5* mRNA reach the levels of wild-type mice, even exceeding them (Fig. 6B, Supplementary Fig. 3A), suggesting a wider recovery of CDKL5 expression in this brain region.

**Figure 6.**
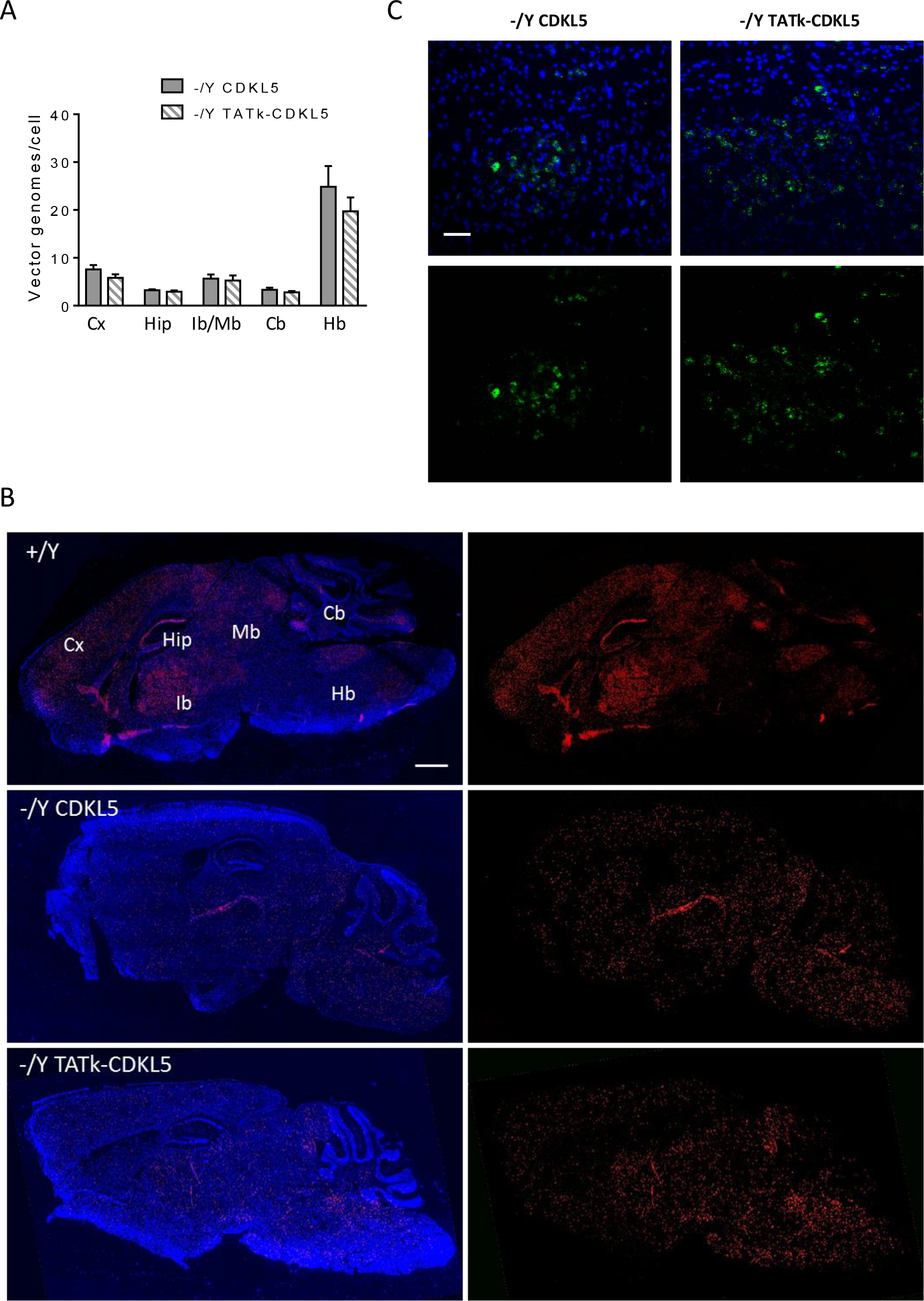
AAV vector transduction and expression and CDKL5 and TATk-CDKL5 proteins in the CNS. (A) Vector genome copy numbers per diploid genomic equivalent in the cortex (Cx, n = 24), hippocampus (Hip, n = 16-15), midbrain-interbrain (Ib-Mb, n = 16-14), cerebellum (Cb, n = 23-24), and hindbrain (Hb, n = 10-9) of *Cdkl5* −/Y mice treated with AAVPHP.B_CDKL5 or AAVPHP.B_Igk-TATk-CDKL5 according to the treatment schedule shown in Supplementary Figure 1A. (B) Representative sagittal brain sections with fluorescence *In Situ* Hybridization (ISH) for *CDKL5* mRNA (red) and nuclei counterstained with DAPI (blue) of a wild-type mouse (+/Y) and of an AAVPHP.B_CDKL5 or AAVPHP.B_Igk-TATk-CDKL5 treated *Cdkl5* −/Y mouse. Scale bar = 1000 μm. (C) Images show CDKL5 and TATk-CDKL5 proteins (green) expression in the hindbrain region of a treated mouse 90 days post-injection. CDKL5 and TATk-CDKL5 proteins expression was evaluated through immunohistochemistry using an anti-HA antibody; nuclei were counterstained with DAPI. Scale bar = 50 µm. Values are presented as means ± SE.

The low level of CDKL5 expression in the brains of treated *Cdkl5* −/Y mice was confirmed through immunohistochemistry (Fig. 6C, Supplementary Fig. 3B) and western blot analysis (Supplementary Fig. 3C) and was reflected by the slight increase in the phosphorylation levels of the direct CDKL5 target, EB2 (Supplementary Fig. 3D,E).

## DISCUSSION

Many chronic neurological diseases do not respond to small molecule therapeutics, and have no effective therapy. Accordingly, no therapies are presently available for the improvement of the neurological phenotypes associated with CDKL5 deficiency disorder. Gene therapy offers the promise of an effective cure for both genetic and acquired brain disease. However, delivering genetic material efficiently to the CNS still remains a hurdle when developing efficacious gene therapy strategies for CNS disorders characterized by widespread neuropathology in several brain regions. Our study provides novel evidence that a gene therapy approach based on a vector carrying the Igk-TATk-CDKL5 transgene provides an increased protein biodistribution and therapeutic efficacy compared to the same vector carrying only CDKL5. We demonstrated that *Cdkl5* −/Y mice treated with AAVPHP.B_Igk-TATk-CDKL5 vector underwent a higher neurodevelopmental and behavioral improvement than mice treated with AAVPHP.B_CDKL5 vector. Importantly, no toxic effects, including immunogenicity problems related to the secreted TATk-CDKL5 protein, were observed in AAVPHP.B_Igk-TATk-CDKL5 treated mice, indicating the safety of this approach. These promising results suggest that a gene therapy with the Igk-TATk-CDKL5 transgene may be an effective approach to treat CDD.

Following AAVPHP.B_CDKL5 vector delivery, *Cdkl5* −/Y mice did not exhibit improvement in behavior in comparison with vehicle-treated mice. The poor therapeutic effect of a gene therapy with only CDKL5 has previously been reported ^31^ and was attributed to the necessity of a more robust brain transduction to ameliorate behavioral deficits in this mouse model ^32^. Noteworthy, we found that the secretable TATk-CDKL5 protein amplifies the therapeutic effect compared to the CDKL5 alone. With the same infection efficacy as that of the AAVPHP.B_CDKL5 vector, a gene therapy with AAVPHP.B_Igk-TATk-CDKL5 was sufficient to improve various behavioral defects in the *Cdkl5* −/Y mouse, such as innate behaviors and motor performance, and ameliorate visual function. Albeit in a more marginal way, hippocampal-dependent learning was improved. The lesser therapeutic effect may be attributed to the lower number of viral copies reaching the hippocampus compared to other brain regions. The correlation between levels of CDKL5 re-expression and effectiveness of the gene therapy was confirmed by the finding that in the brainstem, where the CDKL5 endogenous levels are lower than in the rest of the brain ^33^, treatment with either AAVPHP.B_CDKL5 or AAVPHP.B_Igk-TATk-CDKL5 vectors led to an higher CDKL5 re-expression than in the other brain regions, as suggested by the mRNA level and protein expression. Accordingly, it is suggestive to note how the centers that regulate the appearance of REM sleep are mainly located in the brainstem. The brainstem high re-expression of CDKL5, obtained in *Cdkl5* −/Y mice by treatment with either AAVPHP.B_CDKL5 or AAVPHP.B_Igk-TATk-CDKL5 vectors, could then explain the normalization of the breathing pattern during REM sleep.

As an anatomical substrate of the ameliorated behavioral performance, in AAVPHP.B_Igk-TATk-CDKL5-treated *Cdkl5* −/Y mice we found that their impaired dendritic and synaptic development was restored, as was neuronal survival in the hippocampus. In contrast, the lack of behavioral improvement in *Cdkl5* −/Y mice treated with AAVPHP.B_CDKL5 may be accounted for by the reduced effect of treatment on neuronal survival and dendritic development. Similarly, microglia over-activation, a recently described alteration in the brains of *Cdkl5* −/Y mice ^30^, is inhibited only by treatment with AAVPHP.B_Igk-TATk-CDKL5, further supporting the amplified therapeutic effect of the secretable TATk-CDKL5 protein. The reason why the restoration of several anatomical defects in the hippocampus of AAVPHP.B_Igk-TATk-CDKL5-treated *Cdkl5* −/Y mice does not induce the full recovery of hippocampus-dependent cognitive abilities may be attributable to the complex and intricate *in vivo* brain function which goes beyond structural restorations.

Interestingly, despite finding a broad distribution of the delivered cassettes over the different brain areas of the viral-injected mice, the viral transcript biodistribution differs to that of the wild-types, in which *CDKL5* mRNA is mainly expressed in the cerebral cortex and in the hippocampus. This different distribution could explain the modest therapeutic efficacy showed by the CDKL5 gene therapy. Moreover, we observed unexpected inconsistency between the expression of the viral mRNA and the low CDKL5 protein expression, suggesting a relatively low half-life or reduced translation of the delivered CDKL5. The hypothesis of CDKL5 instability is in agreement with a recent finding showing that the viral *MeCP2* mRNA was not actively translated by ribosomes, underscoring the importance of complex endogenous regulatory elements for MeCP2 protein expression ^34^. Similarly, we found that, while the viral *CDKL5* mRNA levels suggest a recovery of CDKL5 expression in the brain, i.e., in the brainstem, the *CDKL5* expression was much lower compared to that found in wild-type mice, indicating that CDKL5 protein levels correlate poorly with viral mRNA levels. Future studies are needed to characterize regulatory elements of the *CDKL5* mRNA and to design a CDKL5 transgene cassette that would be useful to obtain higher CDKL5 protein levels. However, it was demonstrated that even a modest 5-10% re-expression of MeCP2 has a promising therapeutic effect on the RTT phenotype in a mouse model of RTT ^35^. Although the actual quantity of CDKL5 required to achieve therapeutic efficacy is not known, we recently demonstrated, using a protein substitution therapy approach with a TATk-CDKL5 fusion protein ^17^, that the amount of CDKL5 protein necessary to rescue neurological phenotypes of a mouse model of CDD is very small ^17^. Here we confirmed that the low CDKL5 levels re-expressed in the brains of AAV vector-treated *Cdkl5* −/Y mice are sufficient to improve CDD phenotypes if supported by an increased biodistribution due to the properties of the Igk-TATk fusion protein.

This study has provided a first proof-of-principle that an innovative gene therapy approach based on the unique advantages of the Igk-TATk-CDKL5 transgene is highly efficient in improving neurodevelopmental and behavioral impairments in a mouse model of CDD. Such promising results imply that this approach may become a powerful tool for the cure of CDD, and could open avenues to the development of gene therapy for other monogenic diseases based on the unique and compelling properties of the Igk-TATk-fusion protein approach.

## MATERIALS AND METHODS

### Cloning of viral plasmids and production of AAV vectors

To express the Igk-TATk-CDKL5 and CDKL5 proteins by means of AAV vector-mediated gene delivery, the Igk-TATk-hCDKL51 or hCDKL51 gene expression cassettes were subcloned in the backbone of pAAV-CBh-DIO-EGFP (plasmid #87168, Addgene). The designed viral cassettes between the two AAV2 inverted terminal repeats (ITRs) contain a CBh promoter (0.8 kbp), the Igk-TATk-CDKL5 (3.1 kbp) or CDKL51 (2.9 kbp) open reading frame, followed by a WPRE and an SV40 polyadenylation signal (0.4 kbp). The TATk-CDKL5 and CDKL5 proteins were tagged with a haemagglutinin (HA)-tag. The AAV vector genomes containing the above-described expression cassettes were packaged in the AAVPHP.B capsid in HEK293 cells using an adenovirus-free triple transfection method (for details, see Supplementary material).

### Cell lines and primary cultures

HEK293T cell line was maintained in Dulbecco Modified Eagle Medium (DMEM, Gibco) supplemented with 10% heat-inactivated FBS (Gibco), 2 mM of glutamine (Gibco), and antibiotics (penicillin, 100 U/mL; streptomycin, 100 μg/mL; Gibco), in a humidified atmosphere of 5% of CO2 at 37°C. Cell medium was replaced every 3 days and the cells were sub-cultured once they reached 90% confluence. Cells were transfected with designated plasmid DNA using Metafectene Easy Plus (Biontex). Forty-eight hours after transfection, cells were harvested, washed in PBS, and lysed for total protein extraction. Cell medium was also collected and 200x concentrated as described previously ^17^. Cell extracts and medium were used for western blot analysis.

Primary hippocampal neuronal cultures were prepared from 1-day-old (P1) wild-type and *Cdkl5* −/Y mice as previously described ^36^. Briefly, hippocampi were dissected from mouse brains under a dissection microscope and treated with trypsin (Gibco) for 15 min at 37°C and DNase I (Sigma-Aldrich) for 2 min at room temperature before being triturated mechanically with a fire-polished glass pipette to obtain a single-cell suspension. Cells were plated on coverslips coated with poly-L-lysine in 6-well plates and cultured in Neurobasal medium (Gibco) supplemented with B27 (Invitrogen) and glutamine (Gibco). Cells were maintained in vitro at 37°C in a 5% CO2-humified incubator.

### *In vitro* AAV transduction

Primary hippocampal neurons were infected with AAVPHP.B_Igk-TATk-CDKL5 and AAVPHP.B_CDKL5 (MOI of 10^6^) at day in vitro (DIV) 2, fixed at DIV7 with 4% paraformaldehyde + 4% sucrose in 100 mM phosphate buffer pH 7.4. Fixed cells were stained with the primary and secondary antibodies listed in Supplementary Table 1. Nuclei were counterstained with Hoechst-33342 (Sigma-Aldrich) and fluorescent images were acquired using a Nikon Eclipse Te600 microscope equipped with a Nikon Digital Camera DXM1200 ATI system (Nikon Instruments, Inc. Melville, NY, USA).

### Co-culture system

For co-culture experiments, HEK293T cells were plated in a 6-well plate, while primary hippocampal neurons were plated on cover glasses. Twenty-four hours after plating, HEK293T cells were transfected with the AAV vector plasmid containing the Igk-TATk-CDKL5 or CDKL5 cassette. Twenty-four hours after transfection, the co-culture was prepared as follows: HEK293T cells were washed twice with fresh neuronal culture medium. The cover glasses with primary hippocampal neurons (DIV5) were transferred to the HEK293T 6-well plate in an elevated position of the gel supports underneath. After 48 h of co-culturing, the cover glasses with neurons were removed; neurons were washed in PBS, fixed with 4% paraformaldehyde + 4% sucrose in 100 mM phosphate buffer pH 7.4 and processed for immunocytochemistry. The primary and secondary antibodies used are listed in Supplementary Table 1.

### Animal husbandry

The mice used in this work derive from the *Cdkl5* −/Y strain in the C57BL/6N background developed in ^8^ and backcrossed in C57BL/6J for three generations. Animals were genotyped as previously described ^8^. Age-matched wild-type (+/Y) littermates were used for all experiments. The day of birth was designated as postnatal day (P) zero and animals with 24 h of age were considered as 1-day-old animals (P1). Mice were housed 3–5 per cage on a 12 h light/dark cycle in a temperature-(23 °C) and humidity-controlled environment with standard mouse chow and water *ad libitum*. The animals’ health and comfort were controlled by the veterinary service. All research and animal care procedures were performed in accordance with the Italian and European Community law for the use of experimental animals and were approved by Bologna University Bioethical Committee. All efforts were made to minimize animal suffering and to keep the number of animals used to a minimum.

### In vivo AAV delivery

*Intraventricular infusion –* Neonatal injections were carried out as previously described ^37^. 10^11^ viral vector genomes were injected into 1-day post-gestation (P1) neonatal *Cdkl5* −/Y mice via ICV injection targeting the anterior horn of the lateral ventricle. Prior to the procedure pups were incubated on ice for 1 min and subsequently injected using a 33-gauge needle (Hamilton, Reno, NV, USA). Injected neonates were subsequently returned to the dam. Mice were sacrificed 2 months (P60) post-injection.

*Intracarotid infusion* - Surgery was performed under general anesthesia (ISOFLO, Esteve Spa, 1.8-2.4% in oxygen, inhalation route) with the mouse’s body temperature maintained at 37°C by using a heating pad and intra-operative analgesia (10 µL of Norocarp dissolved in 1 ml of saline; 0.2 ml subcutaneously, Pfizer). All procedures were performed in sterile conditions. The tip of the catheter was flushed with sterile heparin. The isolation of the carotid artery was performed as previously described ^38^. Once isolated, two silk suture threads (Softsilk 5-0) were proximally and distally placed below the arterial segment of interest. The proximal thread (posterior, close to the heart) was permanently knotted and tied while the distal one (near the bifurcation of the carotid) was only pulled by another operator to temporarily close off the artery and to keep the artery slightly taut during the insertion of the catheter. The catheter was inserted between the suture threads, close to the proximal thread, through a hole made by a 90° bent needle (25 G). After the tip of the catheter was inserted, the viral solution was infused with 50 µl/min speed, by using an infusion pump (Harvard Apparatus, Holliston, MA, USA). Mice were injected with a dose of 10^12^ vg/mouse. When all the amount of the solution (200ul) was infused, the infusion was stopped and the catheter was gently pulled out. The threads were both tied. Finally, suture stitches and an antiseptic ointment (Betadine 10%, Viatris) were applied to the skin incision. At the end of the surgical procedure, an antibiotic solution (30 μL of Veterinary Rubrocillin, Intervet, Schering-Plow Animal Health, dissolved in 0.8 ml of sterile saline) was administered subcutaneously to prevent infections and to rehydrate the animal.

### Behavioral Testing

Behavioral tests were performed 2 months after the intracarotid infusion. The sequence of the tests was arranged to minimize the possibility of one test influencing the subsequent evaluation of the next test, and mice were allowed to recover for 2 days between different tests. Mice were allowed to habituate to the testing room for at least 1 h before the test, and testing was always performed at the same time of day. Behavioral studies were carried out on the saline-injected *Cdkl5* +/Y and *Cdkl5* −/Y mice groups and AAV-injected *Cdkl5* −/Y mice groups. A total of 87 animals divided into three independent test cohorts were used. The first test cohort consisted of 34 animals (*Cdkl5* +/Y + vehicle n = 10, *Cdkl5* −/Y + vehicle n = 6, *Cdkl5* −/Y + AAVPHP.B_CDKL5 n = 10, and *Cdkl5* −/Y + AAVPHP.B_Igk-TATk-CDKL5 n = 8) that were tested with the following assays: marble burying, nesting test, hindlimb clasping, open field, Barnes maze. The second cohort consisted of 34 animals (*Cdkl5* +/Y + vehicle n = 4, *Cdkl5* −/Y + vehicle n = 10, *Cdkl5* −/Y + AAVPHP.B_CDKL5 n = 10, and *Cdkl5* −/Y + AAVPHP.B_Igk-TATk-CDKL5 n = 10) that were tested with the following assays: marble burying, nesting test, hindlimb clasping, open field. The third cohort consisted of 19 animals (*Cdkl5* +/Y + vehicle n = 6, *Cdkl5* −/Y + AAVPHP.B_CDKL5 n = 6, and *Cdkl5* −/Y + AAVPHP.B_Igk-TATk-CDKL5 n = 7) that were tested with the following assays: marble burying, nesting test, hindlimb clasping, open field, Barnes maze. See Supplementary material for detailed behavioral methods.

The overall behavioral improvement of *Cdkl*5 −/Y mice subjected to gene therapy with AAVPHP.B_Igk-TATk-CDKL5 or AAVPHP.B_CDKL5 vector was evaluated through an average behavioral score for each genotype and treatment in different tests. Mice were assessed on a 1-4 scale for each behavioral test: marble burying, nesting test, hindlimb clasping, open field (velocity and distance moved), and Barnes maze tests. For each test, the gap between the minimum and maximum values was then divided into four ranges. Scores of 1–4 for each quartile were assigned based on performance increase. We included the 5-to 15-minute interval for the open field test (velocity and distance moved) in the global score evaluation.

### Non-invasive assessment of sleep and breathing pattern

Hypnic and respiratory phenotypes of mice were assessed non-invasively with a validated technique based on whole-body plethysmography (WBP) ^22,39^. Briefly, 2 months after the intracarotid injection each mouse was placed inside a WBP chamber flushed with air at 1.5 l/h for the first 8 h of the light period. The respiratory (WBP chamber pressure) signal was continuously recorded together with chamber humidity and temperature, digitized, and stored at 128 Hz, 4 Hz, and 4 Hz, respectively. The system was calibrated with a 100 μL micro-syringe (Hamilton, Reno, USA) at the end of each recording. The states of wakefulness, non-rapid-eye-movement sleep (NREMS), and rapid-eye-movement sleep (REMS) were scored based on inspection of the raw WBP signal with the investigators blind to the animal’s genotype. Quantitative analysis of breathing was restricted to stable sleep episodes ≥ 12 s because of the frequent occurrence of movement artefacts during wakefulness. Apneas were automatically detected as breaths with instantaneous total breath duration (TTOT) > 3 times; the average TTOT for each mouse and sleep state, and detection accuracy were checked on raw recordings.

### Assessment of visual responses

The methods employed in ^23^ were used. In anesthetized mice the scalp was removed, and the skull carefully cleaned with saline. The skin was secured to the skull using cyanoacrylate. Then a thin layer of cyanoacrylate was poured over the exposed skull to attach a custom-made metal ring (9 mm internal diameter) centered over the binocular visual cortex. After surgery, the animals were left to recover for at least 1 week and then injected intracarotidally with a dose of 10^12^ vg/mouse. Non-invasive transcranial intrinsic optical signal (IOS) recordings were performed 45 days later under isoflurane anesthesia (0.5-1%), supplemented with an intraperitoneal injection of chlorprothixene hydrochloride (1.25 mg/kg). Images were obtained using an Olympus microscope (BX50WI). Red light illumination was provided by 8 red LEDs (625 nm, Knight Lites KSB1385-1P) attached to the objective (Zeiss Plan-NEOFLUAR 5x, NA: 0.16) using a custom-made metal LED holder. Visual evoked responses were quantitatively measured as reported in ^23^.

### CDKL5 mRNA and proteins detection

Mice were perfused with 4% paraformaldehyde in 100 mM phosphate buffer (pH 7.4). Brains were collected and cut along the midline. Hemispheres were submerged in 4% paraformaldehyde in 100 mM phosphate buffer (pH 7.4) for 24h at 4°C and then let sink in sucrose 15%, before being frozen at −80°C. The hemispheres were then cut with a cryostat into 15 um-thick sagittal sections which were serially collected on glasses. The *In Situ* Hybridization (ISH) for the *CDKL5* RNA was performed with the Base Scope® technology (Biotechne) following the manufacturer’s protocol using a 1ZZ probe designed on the CDKL5 exon 4. For double staining, the ISH was followed by immunohistochemistry for CDKL5 proteins detection.

For immunohistochemistry, brain sections were incubated overnight at 4°C with a primary anti-HA antibody (Supplementary Table 1) and for 2 h with an HRP-conjugated anti-rabbit secondary antibody (Supplementary Table 1). Detection was performed using the either the TSA Cyanine 3 Plus or the TSA Plus Fluorescein Evaluation Kits (Perkin Elmer).

### Immunohistochemistry procedures

Mice were sedated with isoflurane (2% pure oxygen) and sacrificed for cervical dislocation. The brains were quickly removed and cut along the midline. The two hemispheres were fixed separately by immersion in a solution of 4% paraformaldehyde in 100 mM phosphate buffer (pH 7.4) for 48 h and subsequently stored in 20% sucrose for another 24 h, before being frozen in dry ice and kept at −80 °C. The hemispheres were then cut with a freezer microtome into 30 μm-thick coronal sections which were serially collected in a 96-well plate containing a solution consisting of 30% glycerol, 30% ethylene glycol, 0.02% sodium azide in 1X PBS. The primary and secondary antibodies used are listed in Supplementary Table 1.

For quantification of PSD-95 immunoreactive puncta, images from the CA1 layer were acquired using a LEICA TCS SL confocal microscope (LEITZ; Leica Microsystems, Wetzlar, Germany; objective 63X, NA 1.32; zoom factor = 8). Three to four sections per animal were analysed and the number of PSD-95 immunoreactive puncta was expressed per μm^2^.

### Golgi staining, neuronal tracing and spine evaluation

Hemispheres were Golgi-stained using the FD Rapid GolgiStain TM Kit (FD NeuroTechnologies) as previously described ^40^. Dendritic trees of Golgi-stained apical dendritic branches of CA1 field neurons were traced with a dedicated software that was custom-designed for dendritic reconstruction (Immagini Computer), interfaced with Image Pro Plus (Media Cybernetics). Dendritic spine density was measured by manually counting the number of dendritic spines. In each mouse, 10-15 dendritic segments (segment length: 10 µm) from each zone were analysed and spine density was expressed as the total number of spines per 10 µm. Based on their morphology, dendritic spines can be divided into two different categories that reflect their state of maturation: immature spines and mature spines. The number of spines belonging to each class was counted and expressed as a percentage.

### Western Blotting

HEK293T cells transfected with plasmid DNA were lysates in Laemmli buffer supplemented with β- mercaptoethanol, sonicated and boiled at 95 °C for 10 min. Brains of treated *Cdkl5* −/Y mice were homogenized in ice-cold RIPA buffer supplemented with 1mM PMSF, and with 1% protease and phosphatase inhibitor cocktail (Sigma-Aldrich). Equivalent amounts of protein were subjected to electrophoresis on a 4-12% Mini-PROTEAN® TGXTM Gel (Bio-Rad) and transferred to a Hybond-ECL nitrocellulose membrane (Amersham - GE Healthcare Life Sciences). The primary and secondary antibodies used are listed in Supplementary Table 1.

### Real-Time PCR

Animals were sedated with isoflurane (2% pure oxygen) and sacrificed for cervical dislocation. The brain was quickly removed and the various brain regions were dissected and stored at −80 °C.

*Viral biodistribution analysis* - Genomic DNA (gDNA) was extracted from the brain region of interest with the NucleoSpin®Tissue Kit (Macherey-Nagel) extraction kit and 100 ng of gDNA was used as a qPCR template. For the quantification of the number of viral copies, a portion of the viral promoter CBh was amplified (Fw 5’-TACTCCCACAGGTGAGCGG-3’, Rev 5’-GGCAGGTGCTCCAGGTAAT-3’). For data normalization a portion of the mAgouti gene was also amplified (Fw 5’-GGCGTGGTCAGTGGTTGTG-3 ‘, Rev 5’-TTTAGCTTCCACTAGGTTTCCTAGAAA-3’). For data interpolation, the calibration curves for CBh and mAgouti were generated through the quantification of serial dilutions of the AAV vector plasmid and a plasmid containing Agouti fragment, respectively. A master calibration curve for CBh and mAgouti was calculated as the mean of several calibration curves obtained from multiple runs (N = 9, for CBh; N = 12, for mAgouti) ^41^. Ct values were interpolated in the master calibration curve to obtain the number of viral copies for each sample and the mAgouti copy number for internal normalization. Finally, viral copy number/cell was calculated for each sample considering the murine genomic DNA molecular weight of 3pg (6pg per diploid cells).

*Analysis of AAV vector genome transcripts* - Total RNA was isolated from the brains of *Cdkl5* −/Y mice treated with vehicle, AAVPHP.B_CDKL5, or AAVPHP.B_Igk-TATk-CDKL5 with the GRS FullSample Purification Kit (GRISP) according to the manufacturer’s instruction. Isolated mRNA was subjected to a DNase I treatment (GRISP), and cDNA synthesis was achieved using iScript™ Advanced cDNA Synthesis Kit (Bio-Rad), according to the manufacturer’s instruction. Reverse transcriptase-PCRs were performed using SsoAdvanced Universal SYBR Green Supermix (Bio-Rad) in iQ5 Real-Time PCR Detection System (Bio-Rad). A portion of the m-hCDKL5 (Fw 5’-CTTAAATGCAGACACAAGGAAACAC-3 ‘, Rev 5’-CGAAGCATTTTAAGCTCTCGT-3’) sequence was amplified for the quantification of AAV vector genome transcripts. A portion of mGAPDH (Fw 5’-GAACATCATCCCTGCATCCA-3’, Rev 5’-CCAGTGAGCTTCCCGTTCA-3’) sequence was amplified for data normalization. The differential folds of expression were calculated using the ΔΔCt method. Values were expressed as the fold increase in the CDKL5 expression in the cortex relative to that of the wild types.

### Statistical Analysis

Results are presented as mean ± standard error of the mean (± SE), and n indicates the number of mice. Statistical analysis was performed using GraphPad Prism software (GraphPad Software, Inc., San Diego, CA). All datasets were analysed using the ROUT method (Q=1%) for the identification of significant outliers and the Shapiro-Wilk test for normality testing. Datasets with normal distribution were analysed for significance using Student’s t-test or an ordinary one-way analysis of variance (ordinary one-way ANOVA). *Post hoc* multiple comparisons were carried out using the Fisher least significant difference (Fisher’s LSD) or a Tukey test. Datasets with non-parametric distribution were analysed using the Kruskal-Wallis test. *Post hoc* multiple comparisons were carried out using Dunn’s multiple comparison test or the Mann-Whitney U test. For the open field and the learning phase of the Barnes maze, statistical analysis was performed using a repeated measure two-way analysis of variance (RM two-way ANOVA). A probability level of p<0.05 was considered to be statistically significant.

## AUTHORS CONTRIBUTION

EC, HN, ST, and GM designed the study. GM, MR and GG performed the experiments and analysed the data. LG, ML, NM, ST helped with data collection and analysis. SB, SA, and CB performed the assessment of the breathing pattern and the viral injections. GS and LL performed the assessment of visual responses. HRB produced the viruses. GZ, MG, AM, HN, TP and ST contributed to the critical revision of the article. EC wrote the manuscript with input from all authors. All authors reviewed the results and approved the final version of the manuscript.

## FUNDING

This work was supported by the Telethon Foundation (grant number GGP19045 to EC and MG), by the Jérôme Lejeune Foundation, the University of Pennsylvania Orphan Disease Center (on behalf of LouLou Foundation), the Italian parent association “CDKL5 insieme verso la cura” (grant to EC and HN) and “CDKL5 Associazione di Volontariato Onlus” (grant to EC), the University of Bologna (“Proof of Concept program” grant to EC) and by a Public Health Service grant (grant number R01 NS088399 to HN).

## CONFLICT OF INTEREST STATEMENT

The authors have no conflicts of interest to declare.

## Notes

### Competing Interest Statement

The authors have declared no competing interest.

## BIBLIOGRAPHY

1 Mari, F. et al. CDKL5 belongs to the same molecular pathway of MeCP2 and it is responsible for the early-onset seizure variant of Rett syndrome. Hum Mol Genet 14, 1935–1946, doi:10.1093/hmg/ddi198 (2005).

2 Bertani, I. et al. Functional consequences of mutations in CDKL5, an X-linked gene involved in infantile spasms and mental retardation. J Biol Chem 281, 32048–32056, doi:10.1074/jbc.M606325200 (2006).

3 Fehr, S. et al. The CDKL5 disorder is an independent clinical entity associated with early-onset encephalopathy. Eur J Hum Genet 21, 266–273, doi:10.1038/ejhg.2012.156 (2013).

4 Guerrini, R. & Parrini, E. Epilepsy in Rett syndrome, and CDKL5-and FOXG1-gene-related encephalopathies. Epilepsia 53, 2067–2078, doi:10.1111/j.1528-1167.2012.03656.x (2012).

5 Olson, H. E. et al. Cyclin-Dependent Kinase-Like 5 Deficiency Disorder: Clinical Review. Pediatric neurology 97, 18–25, doi:10.1016/j.pediatrneurol.2019.02.015 (2019).

6 Demarest, S. et al. Severity Assessment in CDKL5 Deficiency Disorder. Pediatr Neurol 97, 38–42, doi:10.1016/j.pediatrneurol.2019.03.017 (2019).

7 Wang, I. T. et al. Loss of CDKL5 disrupts kinome profile and event-related potentials leading to autistic-like phenotypes in mice. Proceedings of the National Academy of Sciences of the United States of America 109, 21516–21521, doi:10.1073/pnas.1216988110 (2012).

8 Amendola, E. et al. Mapping pathological phenotypes in a mouse model of CDKL5 disorder. PloS one 9, e91613, doi:10.1371/journal.pone.0091613 (2014).

9 Okuda, K. et al. CDKL5 controls postsynaptic localization of GluN2B-containing NMDA receptors in the hippocampus and regulates seizure susceptibility. Neurobiol Dis 106, 158–170, doi:10.1016/j.nbd.2017.07.002 (2017).

10 Fuchs, C. et al. Loss of CDKL5 impairs survival and dendritic growth of newborn neurons by altering AKT/GSK-3beta signaling. Neurobiol Dis 70, 53–68, doi:10.1016/j.nbd.2014.06.006 (2014).

11 Fuchs, C. et al. Inhibition of GSK3beta rescues hippocampal development and learning in a mouse model of CDKL5 disorder. Neurobiol Dis 82, 298–310, doi:10.1016/j.nbd.2015.06.018 (2015).

12 Ren, E. et al. Functional and Structural Impairments in the Perirhinal Cortex of a Mouse Model of CDKL5 Deficiency Disorder Are Rescued by a TrkB Agonist. Front Cell Neurosci 13, 169, doi:10.3389/fncel.2019.00169 (2019).

13 Gray, S. J. Gene therapy and neurodevelopmental disorders. Neuropharmacology 68, 136–142, doi:10.1016/j.neuropharm.2012.06.024 (2013).

14 Manno, C. S. et al. Successful transduction of liver in hemophilia by AAV-Factor IX and limitations imposed by the host immune response. Nat Med 12, 342–347, doi:10.1038/nm1358 (2006).

15 Nathwani, A. C. et al. Long-term safety and efficacy of factor IX gene therapy in hemophilia B. N Engl J Med 371, 1994–2004, doi:10.1056/NEJMoa1407309 (2014).

16 Donsante, A. et al. AAV vector integration sites in mouse hepatocellular carcinoma. Science 317, 477, doi:10.1126/science.1142658 (2007).

17 Trazzi, S. et al. CDKL5 protein substitution therapy rescues neurological phenotypes of a mouse model of CDKL5 disorder. Hum Mol Genet 27, 1572–1592, doi:10.1093/hmg/ddy064 (2018).

18 Gray, S. J. et al. Optimizing promoters for recombinant adeno-associated virus-mediated gene expression in the peripheral and central nervous system using self-complementary vectors. Human gene therapy 22, 1143–1153, doi:10.1089/hum.2010.245 (2011).

19 Fuchs, C. et al. Heterozygous CDKL5 Knockout Female Mice Are a Valuable Animal Model for CDKL5 Disorder. Neural Plast 2018, 9726950, doi:10.1155/2018/9726950 (2018).

20 Bahi-Buisson, N. & Bienvenu, T. CDKL5-Related Disorders: From Clinical Description to Molecular Genetics. Mol Syndromol 2, 137–152, doi:000331333 (2012).

21 Okuda, K. et al. Comprehensive behavioral analysis of the Cdkl5 knockout mice revealed significant enhancement in anxiety-and fear-related behaviors and impairment in both acquisition and long-term retention of spatial reference memory. PloS one 13, e0196587, doi:10.1371/journal.pone.0196587 (2018).

22 Lo Martire, V. et al. CDKL5 deficiency entails sleep apneas in mice. J Sleep Res 26, 495–497, doi:10.1111/jsr.12512 (2017).

23 Mazziotti, R. et al. Searching for biomarkers of CDKL5 disorder: early-onset visual impairment in CDKL5 mutant mice. Hum Mol Genet 26, 2290–2298, doi:10.1093/hmg/ddx119 (2017).

24 Fuchs, C. et al. Treatment with the GSK3-beta inhibitor Tideglusib improves hippocampal development and memory performance in juvenile, but not adult, Cdkl5 knockout mice. Eur J Neurosci 47, 1054–1066, doi:10.1111/ejn.13923 (2018).

25 Trazzi, S. et al. HDAC4: a key factor underlying brain developmental alterations in CDKL5 disorder. Hum Mol Genet 25, 3887–3907, doi:10.1093/hmg/ddw231 (2016).

26 Della Sala, G. et al. Dendritic Spine Instability in a Mouse Model of CDKL5 Disorder Is Rescued by Insulin-like Growth Factor 1. Biol Psychiatry 80, 302–311, doi:10.1016/j.biopsych.2015.08.028 (2016).

27 Ricciardi, S. et al. CDKL5 ensures excitatory synapse stability by reinforcing NGL-1-PSD95 interaction in the postsynaptic compartment and is impaired in patient iPSC-derived neurons. Nat Cell Biol 14, 911–923, doi:10.1038/ncb2566 (2012).

28 Loi, M. et al. Treatment with a GSK-3beta/HDAC Dual Inhibitor Restores Neuronal Survival and Maturation in an In Vitro and In Vivo Model of CDKL5 Deficiency Disorder. Int J Mol Sci 22, doi:10.3390/ijms22115950 (2021).

29 Gennaccaro, L. et al. Age-Related Cognitive and Motor Decline in a Mouse Model of CDKL5 Deficiency Disorder is Associated with Increased Neuronal Senescence and Death. Aging and disease (2021).

30 Galvani, G. et al. Inhibition of Microglia Over-activation Restores Neuronal Survival in a Mouse Model of CDKL5 Deficent Disorder. Journal of Neuroinflammation Preprints, doi:doi.org/10.21203/rs.3.rs-203260/v1 (2021).

31 Gao, Y. et al. Gene replacement ameliorates deficits in mouse and human models of cyclin-dependent kinase-like 5 disorder. Brain : a journal of neurology 143, 811–832, doi:10.1093/brain/awaa028 (2020).

32 Benke, T. A. & Kind, P. C. Proof-of-concept for a gene replacement approach to CDKL5 deficiency disorder. Brain : a journal of neurology 143, 716–718, doi:10.1093/brain/awaa055 (2020).

33 Kilstrup-Nielsen, C. et al. What we know and would like to know about CDKL5 and its involvement in epileptic encephalopathy. Neural Plast 2012, 728267, doi:10.1155/2012/728267 (2012).

34 Luoni, M. et al. Whole brain delivery of an instability-prone Mecp2 transgene improves behavioral and molecular pathological defects in mouse models of Rett syndrome. Elife 9, doi:10.7554/eLife.52629 (2020).

35 Carrette, L. L. G., Blum, R., Ma, W., Kelleher, R. J., 3rd & Lee, J. T. Tsix-Mecp2 female mouse model for Rett syndrome reveals that low-level MECP2 expression extends life and improves neuromotor function. Proceedings of the National Academy of Sciences of the United States of America 115, 8185–8190, doi:10.1073/pnas.1800931115 (2018).

36 Beaudoin, G. M., 3rd et al. Culturing pyramidal neurons from the early postnatal mouse hippocampus and cortex. Nat Protoc 7, 1741–1754, doi:10.1038/nprot.2012.099 (2012).

37 Kim, J. Y., Grunke, S. D., Levites, Y., Golde, T. E. & Jankowsky, J. L. Intracerebroventricular viral injection of the neonatal mouse brain for persistent and widespread neuronal transduction. J Vis Exp, 51863, doi:10.3791/51863 (2014).

38 Jacobs, J. D. & Hopper-Borge, E. A. Carotid artery infusions for pharmacokinetic and pharmacodynamic analysis of taxanes in mice. Journal of visualized experiments : JoVE, e51917, doi:10.3791/51917 (2014).

39 Bastianini, S. et al. Accurate discrimination of the wake-sleep states of mice using non-invasive whole-body plethysmography. Scientific reports 7, 41698, doi:10.1038/srep41698 (2017).

40 Guidi, S. et al. Early pharmacotherapy with fluoxetine rescues dendritic pathology in the Ts65Dn mouse model of down syndrome. Brain Pathol 23, 129–143, doi:10.1111/j.1750-3639.2012.00624.x (2013).

41 Sivaganesan, M., Haugland, R. A., Chern, E. C. & Shanks, O. C. Improved strategies and optimization of calibration models for real-time PCR absolute quantification. Water research 44, 4726–4735, doi:10.1016/j.watres.2010.07.066 (2010).

