## Supplementary Figures for "EXPRESSION OF A SECRETABLE, CELL-PENETRATING CDKL5 PROTEIN ENHANCES THE EFFICACY OF AAV VECTOR-MEDIATED GENE THERAPY FOR CDKL5 DEFICIENCY DISORDER"

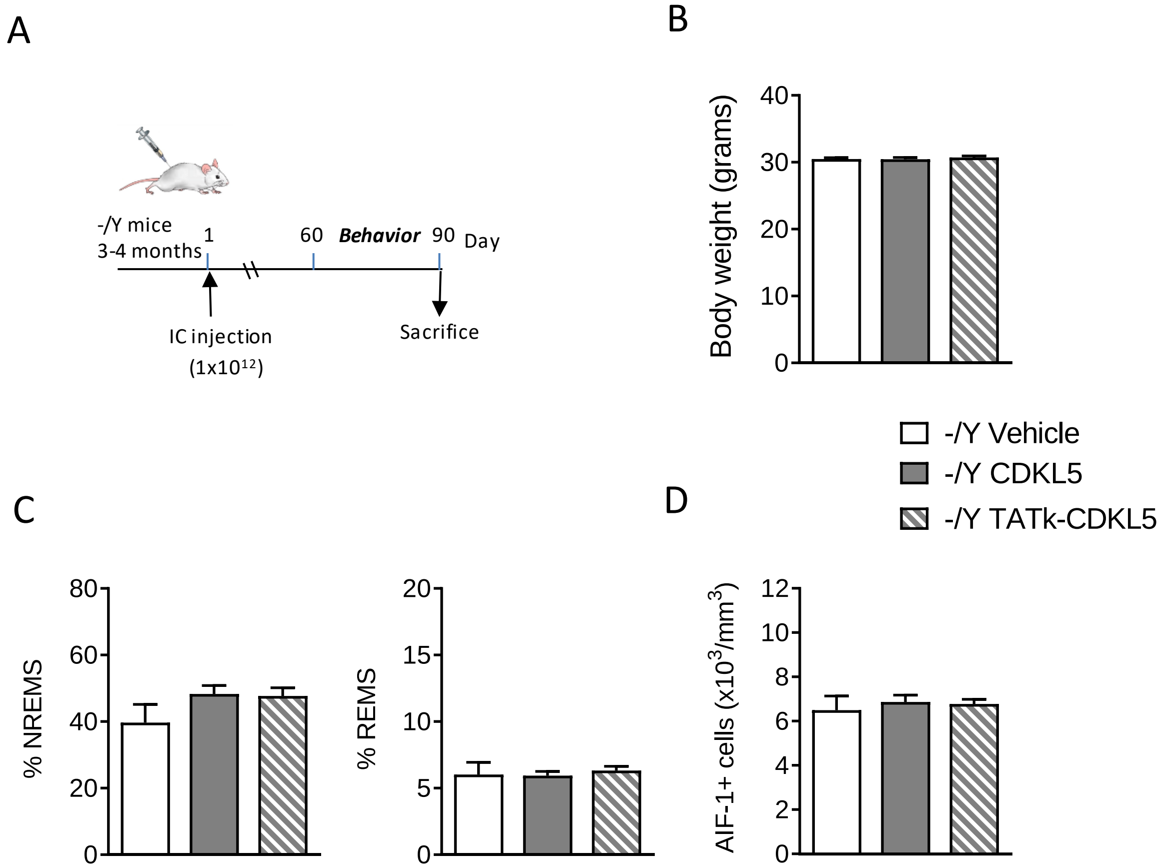


**Supplementary Figure 1**

(A) Experimental design. Adult mice (3-4 months old) were systemically treated (intracarotid injection; IC) with vehicle only, AAVPHP.B_CDKL5, or AAVPHP.B_Igk-TATk-CDKL5, and brain samples were collected 90 days post-injection. (B) Body weight (in grams) of vehicle-treated *Cdkl5* -/Y mice (-/Y; n = 10) and *Cdkl5* -/Y mice treated with AAVPHP.B_CDKL5 (n = 16) or AAVPHP.B_Igk-TATk-CDKL5 (n = 18) vectors according to the treatment schedule shown in A. Mice were weighed before sacrifice at 90 days post-injection. (C) Percentage of time spent in non-rapid-eye-movement sleep (NREMS) and rapid-eye-movement sleep (REMS) during whole-body-plethysmography recordings of vehicle-treated *Cdkl5* -/Y mice (n = 7) and *Cdkl5* -/Y mice treated with AAVPHP.B_CDKL5 (n = 16) or AAVPHP.B_Igk-TATk-CDKL5 (n = 16) vectors. (D) Quantification of AIF-1-positive cells in the hippocampus of vehicle-treated *Cdkl5* -/Y mice (n = 3) and *Cdkl5* -/Y mice treated with AAVPHP.B_CDKL5 (n = 4) or AAVPHP.B_Igk-TATk-CDKL5 (n = 4) vectors. Values are presented as means ± SE.

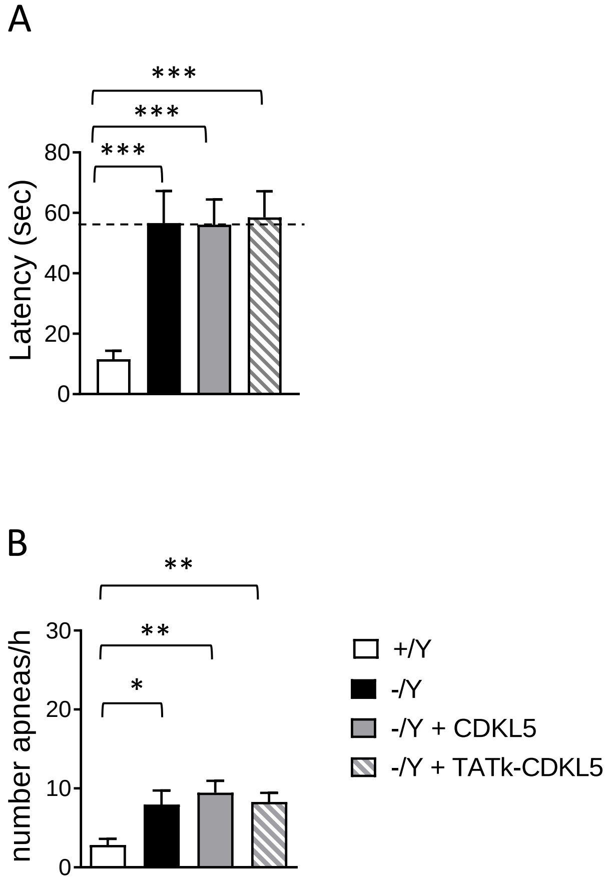


**Supplementary Figure 2**

(A) Spatial memory assessed using the Barnes Maze in wild-type mice (+/Y, n = 16) and *Cdkl5* -/Y mice (n = 10), and in *Cdkl5* -/Y mice treated with AAVPHP.B_CDKL5 (n = 15) or AAVPHP.B_Igk-TATk-CDKL5 (n = 13). The graph shows the latency to find the target hole on the probe day (day 4). (B) Sleep apnea occurrence rate in treated *Cdkl5* -/Y mice was assessed using whole-body plethysmography. Sleep apnea occurrence in vehicle-treated wild-type (+/Y, n = 9) and *Cdkl5* -/Y (n = 7) mice, and in *Cdkl5* -/Y mice treated with AAVPHP.B_CDKL5 (n = 16) or AAVPHP.B_Igk-TATk-CDKL5 (n = 15), during non-rapid eye movement sleep (NREMS). Values are presented as means ± SE. *p < 0.05; **p < 0.01; ***p < 0.001 (Dataset in A, Dunn's test after a Kruskal-Wallis test; dataset in B, Fisher's LSD test after one-way ANOVA).

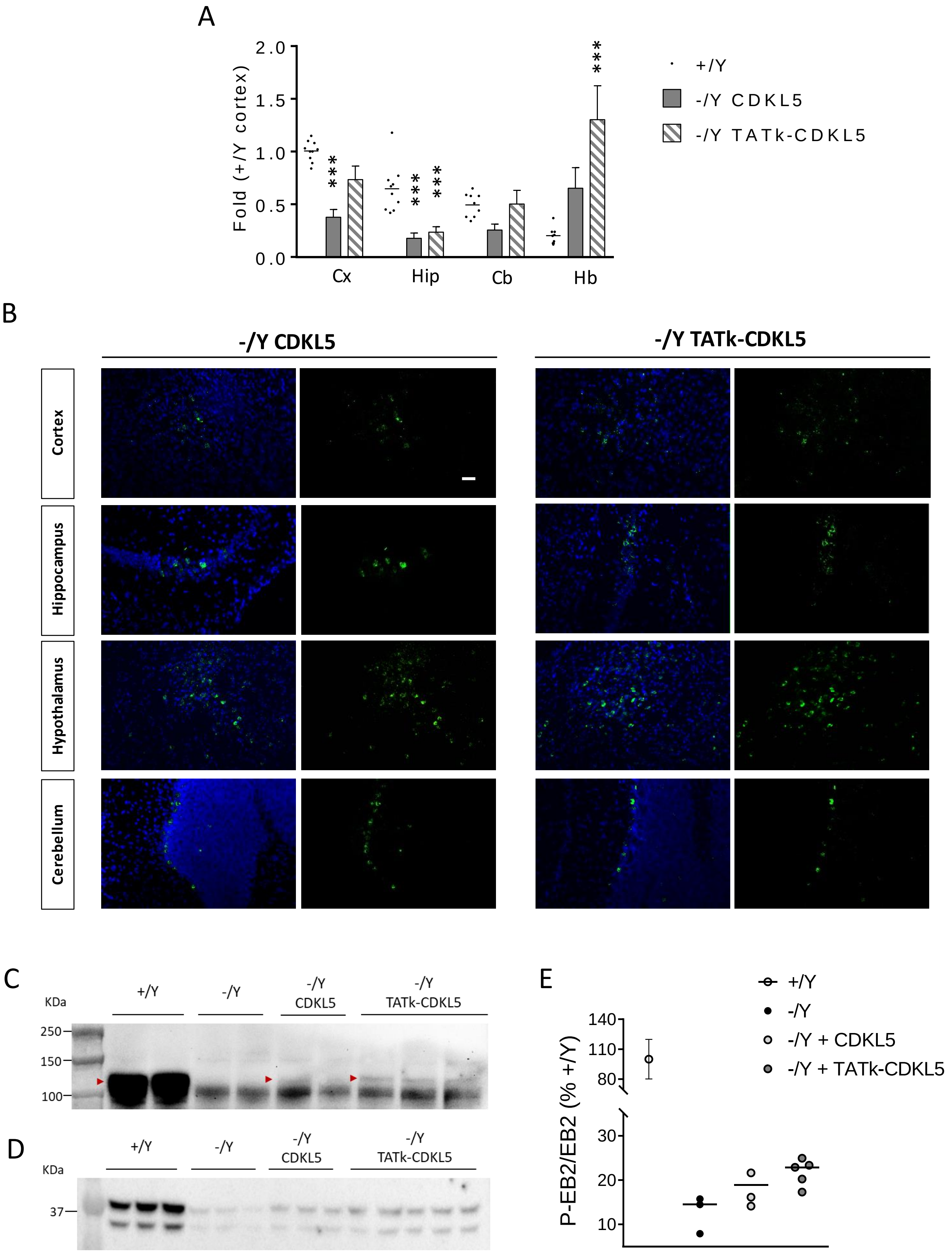

A
Fold (+/Y cortex)
2.0
1.5
1.0
0.5
0.0
Cx
Hip
Cb
Hb
+/Y
-/Y CDKL5
-/Y TATk-CDKL5
B
-/Y CDKL5
-/Y TATk-CDKL5
Cortex
Hippocampus
Hypothalamus
Cerebellum
C
KDa
+/Y
-/Y
-/Y CDKL5
-/Y TATk-CDKL5
250
150
100
D
KDa
37
E
P-EB2/EB2 (% +/Y)
140
110
80
30
20
10
+/Y
-/Y
-/Y + CDKL5
-/Y + TATk-CDKL5

**Supplementary Figure 3**

(A) Expression of *CDKL5* mRNA in the cortex (Cx, n = 10-15-16), hippocampus (Hip, n = 10-10-11), cerebellum (Cb, n = 9-10-10), and hindbrain (Hb, n = 10-14-14) of vehicle-treated wild-type mice (+/Y) and of *Cdkl5* -/Y mice treated with AAVPHP.B vectors (AAVPHP.B_CDKL5 and AAVPHP.B_Igk-TATk-CDKL5). Data are given as the fold change of the +/Y cortex. (B) Images show CDKL5 and TATk-CDKL5 protein (green) expression in the cortex, hippocampus, hypothalamus and cerebellum of a treated mouse 90 days post-injection. CDKL5 and TATk-CDKL5 proteins expression was evaluated through immunohistochemistry using an anti-HA antibody; nuclei were counterstained with DAPI. Scale bar = 50 μm. (C) Western blot analysis of CDKL5 in hindbrain protein extracts from vehicle-treated wild-type (+/Y, n =2) and *Cdkl5* -/Y (n = 2) mice, and those from *Cdkl5* -/Y mice treated with AAVPHP.B_CDKL5 (n = 2) or AAVPHP.B_Igk-TATk-CDKL5 (n = 3) vectors. Red arrowheads indicate mouse Cdkl5 in *Cdkl5* +/Y extracts, human CDKL5, and TATk-CDKL5 in AAVPHP.B_CDKL5 and AAVPHP.B_Igk-TATk-CDKL5 treated *Cdkl5* -/Y mice. (D,E) Western blot analysis of phospho-EB2 in cortical protein extracts from vehicle-treated wild-type (+/Y, n =3) and *Cdkl5* -/Y (n = 3) mice, and those from *Cdkl5* -/Y mice treated with AAVPHP.B_CDKL5 (n = 3) or AAVPHP.B_Igk-TATk-CDKL5 (n = 5) vectors. Data are expressed as a percentage of +/Y and plotted as dots and median (E). Values in (A) are presented as means ± SE. ***p < 0.001 compared to corresponding region of the vehicle-treated wild-type condition (Fisher's LSD test after one-way ANOVA).
