## Supplementary Materials for "EXPRESSION OF A SECRETABLE, CELL-PENETRATING CDKL5 PROTEIN ENHANCES THE EFFICACY OF AAV VECTOR-MEDIATED GENE THERAPY FOR CDKL5 DEFICIENCY DISORDER"

### Production of AAV vectors

AAV vectors were produced in HEK293 cells using an adenovirus-free three plasmid transfection method [1] with modification on a scale of 1 to 2-liter culture. In brief, HEK 293 cells were grown in Dulbecco's modified Eagle's medium (DMEM) supplemented with 10% fetal bovine serum (FBS), L-glutamine, and penicillin-streptomycin. Immediately before plasmid DNA transfection, the culture media were changed to serum-free media. HEK293 cells were transfected with the following three plasmids at a 1:1:1 ratio using the standard polyethyleneimine (PEI) (1 mg/ml) DNA transfection procedure at a DNA:PEI weight ratio of 1:2. The three plasmids used for AAV vector production were: (1) pHELPER (an adenovirus helper plasmid from Agilent), (2) pHLP-AAV-PHP.B (an AAV helper plasmid supplying AAV2 Rep and AAV-PHP.B capsid proteins, constructed based on an AAV9 helper plasmid obtained from James M. Wilson, University of Pennsylvania), and (3) an AAV vector plasmid containing a transgene expression cassette placed between the two AAV2 inverted terminal repeats (ITRs), constructed based on an AAV vector plasmid obtained from Avigen Inc. Five days after transfection, both media and cells were harvested. The harvested media and cells underwent one cycle of freezing and thawing and the cell debris was removed by centrifugation. The culture medium supernatants were made to 8% polyethylene glycol (PEG) 8000 and 0.5 M NaCl, incubated on ice for 3 h, and spun at 10,000 × g for 30 min to precipitate viral particles. The pellets were resuspended in a buffer containing 50 mM Tris-HCl (pH 8.5) and 2 mM MgCl2, treated with Benzonase (EMD Millipore, Darmstadt, Germany) at a concentration of 200 units per mL for 1 h, and subjected to purification by two rounds of cesium chloride (CsCl) density-gradient ultracentrifugation [2], followed by dialysis with phosphate-buffered saline (PBS) with 0.001% Pluronic F68. The final viral preparations were made in PBS/5 % sorbitol/0.001% Pluronic F68 and stored at -80°C until use. The AAV vector titers were determined through a quantitative dot blot assay [3].

### Behavioral Assays

*Marble Burying* - The Marble Burying test is a commonly used test to analyze repetitive and compulsive behaviors in mice [4]. In this study, the protocol used was that described in Thomas' study [5]. In short, 20 marbles are placed in 4 rows of 5 in 4.5 cm of litter inside standard rat cages (20 cm x 45 cm). Subsequently, the mice were positioned at the corner of the cages, opposite the marbles. Animals were given 30 minutes to explore the cages. After 30 minutes, the mice were removed from the cages and the number of "hidden" marbles was counted. A marble is considered hidden if more than 2/3 of its surface is covered by the litter. At least two operators, unaware of the genotype, evaluated the number of hidden marbles and the assigned scores were averaged to avoid individual bias.

*Nesting test* - The ability to build a nest was evaluated as proposed by Deacon [6]. The animals were placed in individual cages with standard bedding, and a piece of absorbent paper (23 cm × 23 cm) was provided. The nests were independently assessed after 24h by two operators using the following scoring system: 0 - no nest, 1 - primitive flat nest (cushion-shaped, consisting of a flat paper handkerchief that slightly raises a mouse above the litter) , 2 - more complex nest (includes biting the piece of paper and deforming it), 3 - neat and complex cup-shaped nest (with shredded paper woven to form the walls of the cup), and 4 - complex nest with hood, with walls which form a ceiling so that the nest becomes a hollow sphere with an opening.

*Clasping of the hind limbs* - The reflex of the animal to bring the hind legs up to the body while it is suspended upside down and held by its tail is typically defined as clasping. This reflex is an indicator that is often used to identify neurological deficits in rodents [7]. For this behavioral test, the animals were suspended by their tail and the extent of hindlimb clasping was observed for 10 seconds. If the hindlimbs are consistently splayed outward, away from the abdomen, it is assigned a score of 0. If one hindlimb is retracted toward the abdomen for more than 50% of the time suspended, it receives a score of 1. If both hindlimbs are partially retracted toward the abdomen for more than 50% of the time suspended, it receives a score of 2. If its hindlimbs are entirely retracted and touching the abdomen for more than 50% of the time suspended, it receives a score of 3 [8].

*Open Field test* - To assess locomotion, the animals were placed in the center of a square arena (50 × 50 cm) and their behavior was monitored for 20 min using a video camera placed in the center of the arena. Distinctive features of locomotor activity, including total distance traveled and average speed of locomotion, were assessed using the EthoVision15XT software (Noldus Information Technology B.V., Netherlands). Test chambers were cleaned with 70% ethanol between tests.

*Barnes maze* - The test was used to measure spatial memory acquisition and retention. An escape box was placed under one of the holes on a circular platform (1 m in diameter) with 20 holes (each hole was 5 cm in diameter) along the perimeter of the platform (Ugo Basile SRL, Italy) that was elevated 60 cm above the floor. Mice were left to locate the escape box starting from a cylindrical circular chamber in the middle of the maze. Mice were trained to find the escape box for 3 days (3 trials per day, max. 3 min per trial) with a 30 min inter-trial interval. To measure the retention of spatial memory a 90 sec probe trial with the escape box removed was performed 24 h after the last trial. Latency to locate the escape box were recorded using a video tracking system (Ethovision 15XT, Noldus Information Technology, The Netherlands). Mice showing freezing behavior for more than two-thirds of the test time were excluded from the analysis.


1 Matsushita, T. *et al.* Adeno-associated virus vectors can be efficiently produced without helper virus. *Gene therapy* **5**, 938-945, doi:10.1038/sj.gt.3300680 (1998).

2 Grimm, D. *et al.* Preclinical in vivo evaluation of pseudotyped adeno-associated virus vectors for liver gene therapy. *Blood* **102**, 2412-2419, doi:10.1182/blood-2003-02-0495 (2003).

3 Powers, J. M., Chang, X. L., Song, Z. & Nakai, H. A Quantitative Dot Blot Assay for AAV Titration and Its Use for Functional Assessment of the Adeno-associated Virus Assembly-activating Proteins. *Journal of visualized experiments : JoVE*, doi:10.3791/56766 (2018).

4 Angoa-Perez, M., Kane, M. J., Briggs, D. I., Francescutti, D. M. & Kuhn, D. M. Marble burying and nestlet shredding as tests of repetitive, compulsive-like behaviors in mice. *Journal of visualized experiments : JoVE*, 50978, doi:10.3791/50978 (2013).

5 Thomas, A. *et al.* Marble burying reflects a repetitive and perseverative behavior more than novelty-induced anxiety. *Psychopharmacology (Berl)* **204**, 361-373, doi:10.1007/s00213-009-1466-y (2009).

6 Deacon, R. M. Assessing nest building in mice. *Nat Protoc* **1**, 1117-1119, doi:10.1038/nprot.2006.170 (2006).

7 Lalonde, R. & Strazielle, C. Brain regions and genes affecting limb-clasping responses. *Brain Res Rev* **67**, 252-259, doi:10.1016/j.brainresrev.2011.02.005 (2011).

8 Guyenet, S. J. *et al.* A simple composite phenotype scoring system for evaluating mouse models of cerebellar ataxia. *Journal of visualized experiments : JoVE*, doi:10.3791/1787 (2010).
