## Supplementary Table for "EXPRESSION OF A SECRETABLE, CELL-PENETRATING CDKL5 PROTEIN ENHANCES THE EFFICACY OF AAV VECTOR-MEDIATED GENE THERAPY FOR CDKL5 DEFICIENCY DISORDER"

| | Target | Description | Dilution | Manufacturer |
|---|---|---|---|---|
| Primary Antibodies | AIF-1 | Rabbit polyclonal | IHC 1:300 | ThermoFisher |
| | CDKL5 | Sheep polyclonal | WB 1:500 | University of Dundee |
| | EB2 | Rabbit polyclonal | WB 1:1000 | Abcam |
| | P-EB2 | Rabbit polyclonal | WB 1:1000 | CovalAb |
| | HA | Rabbit monoclonal | IHC 1:500<br>ICC 1:200<br>WB 1:1000 | Cell signalling Technology |
| | NeuN | Mouse monoclonal | IHC 1:250 | Merck Millipore |
| | PSD95 | Rabbit polyclonal | IHC 1:1000 | Abcam |
| | TubJ | Mouse monoclonal | ICC 1:100 | Sigma Aldrich |
| Secondary Antibodies | Goat anti-Rabbit IgG HRP-conjugated | | WB 1:5000<br>IHC 1:1000 | Jackson ImmunoResearch |
| | Goat anti-Sheep IgG HRP-conjugated | | WB 1:5000 | Jackson ImmunoResearch |
| | Goat anti-mouse IgG FITC-conjugated | | ICC 1:200 | Jackson ImmunoResearch |
| | Goat anti-rabbit IgG FITC-conjugated | | IHC 1:200 | Jackson ImmunoResearch |
| | Goat anti-rabbit IgG Cy3-conjugated | | IHC 1:1000<br>ICC 1:200 | Jackson ImmunoResearch |

**Supplementary Table 1**

List of primary and secondary antibodies. WB: western blotting, IHC: immunohistochemistry, ICC: immunocytochemistry.
